# Effects of systemic oxytocin and beta-3 receptor agonist (CL 316243) treatment on body weight and adiposity in male diet-induced obese rats

**DOI:** 10.1101/2024.09.27.615550

**Authors:** Jared D. Slattery, June R. Rambousek, Edison Tsui, Mackenzie K. Honeycutt, Matvey Goldberg, James L. Graham, Tomasz A. Wietecha, Tami Wolden-Hanson, Amber L. Williams, Kevin D. O’Brien, Peter J. Havel, James E. Blevins

**Affiliations:** VA Puget Sound Health Care System, Office of Research and Development Medical Research Service, Department of Veterans Affairs Medical Center, Seattle, WA 98108, USA; Division of Metabolism, Endocrinology and Nutrition, Department of Medicine, University of Washington School of Medicine, Seattle, WA 98195, USA; Division of Cardiology, Department of Medicine, University of Washington School of Medicine, Seattle, WA 98195, USA; UW Medicine Diabetes Institute, University of Washington School of Medicine, Seattle, WA 98109; Department of Nutrition, University of California, Davis, CA 95616, USA; Department of Molecular Biosciences, School of Veterinary Medicine, University of California, Davis, CA 95616, USA

**Keywords:** Obesity, brown adipose tissue, white adipose tissue, food intake, oxytocin

## Abstract

Previous studies have implicated hindbrain oxytocin (OT) receptors in the control of food intake and brown adipose tissue (BAT) thermogenesis. We recently demonstrated that hindbrain [fourth ventricle (4V)] administration of oxytocin (OT) could be used as an adjunct to drugs that directly target beta-3 adrenergic receptors (β3-AR) to elicit weight loss in diet-induced obese (DIO) rodents. What remains unclear is whether systemic OT can be used as an adjunct with the β3-AR agonist, CL 316243, to increase BAT thermogenesis and elicit weight loss in DIO rats. We hypothesized that systemic OT and β3-AR agonist (CL 316243) treatment would produce an additive effect to reduce body weight and adiposity in DIO rats by decreasing food intake and stimulating BAT thermogenesis. To test this hypothesis, we determined the effects of systemic (subcutaneous) infusions of OT (50 nmol/day) or vehicle (VEH) when combined with daily systemic (intraperitoneal) injections of CL 316243 (0.5 mg/kg) or VEH on body weight, adiposity, food intake and brown adipose tissue temperature (T_IBAT_). OT and CL 316243 monotherapy decreased body weight by 8.0±0.9% (*P*<0.05) and 8.6±0.6% (*P*<0.05), respectively, but OT in combination with CL 316243 produced more substantial weight loss (14.9±1.0%; *P*<0.05) compared to either treatment alone. These effects were associated with decreased adiposity, energy intake and elevated T_IBAT_ during the treatment period. The findings from the current study suggest that the effects of systemic OT and CL 316243 to elicit weight loss are additive and appear to be driven primarily by OT-elicited changes in food intake and CL 316243-elicited increases in BAT thermogenesis.

## Introduction

The obesity epidemic and its associated complications increase the risk for cardiovascular disease (CVD), hypertension, cancer, type 2 diabetes, and COVID-19 [1; 2]. Many of the monotherapies to treat obesity are of limited effectiveness, associated with adverse and/or unwanted side effects (i.e. diarrhea, nausea, vomiting, sleep disturbance and depression) and/or are poorly tolerated. Improvements have been made in monotherapies to treat obesity, particularly within the family of drugs that target the glucagon-like peptide-1 receptor (GLP-1R). Although the FDA recently approved the use of the long-acting and highly effective GLP-1R agonist, semaglutide [3], it can also be associated with mild to moderate gastrointestinal (GI) side effects [3; 4], underscoring the need for continued optimization of existing treatments.

Recent studies suggest that combination therapy (co-administration of different compounds) and monomeric therapy (dual or triple agonists in single molecule) are more effective than monotherapy for prolonged weight loss [5; 6]. Marked weight loss has been reported in long-term (20 weeks to ≥ 1 year) clinical studies in humans treated with the amylin analogue, cagrilintide, and semaglutide (≈ 15.6 to 17.1% of initial body weight [7; 8]) and the FDA-approved drug, Qsymia (topiramate + phentermine) (≈ 10.9% of initial body weight; [9]). Alternatively, the monomeric compound, tirzepatide (Zepbound^TM^), targets both GLP-1R and glucose-dependent insulinotropic polypeptide receptor (GIPR), and was recently reported to elicit 20.9% and 25.3% weight loss in humans with obesity over 72- [10] and 88-week trials [11]. Similarly, a recently described drug conjugate, GLP- 1-MK-801, which serves as both a GLP-1R agonist and an NMDA receptor antagonist, resulted in 23.2% weight loss following a 14-day treatment regimen in DIO mice [12]. In addition, retatrutide, a triple-agonist that targets GIPR, glucagon receptors (GCGR) and GLP-1R was reported to reduce body weight by 24.2% over a 48-week trial [13]. Despite the considerable improvements that have been made with respect to weight loss, these treatments are still associated with adverse GI side effects [10], leading, in some cases, to the discontinuation of the drug in up to 7.1% of participants [10].

While the hypothalamic neuropeptide, oxytocin (OT) is largely associated with reproductive behavior [14], recent studies implicate an important role for OT in the regulation of body weight [15; 16; 17; 18]. Studies to date indicate that OT elicits weight loss, in part, by reducing food intake and increasing lipolysis [19; 20; 21] and energy expenditure [19; 22; 23; 24]. While OT is effective at evoking prolonged weight loss in DIO rodents [20; 23; 24; 25; 26; 27; 28; 29] and nonhuman primates [19], its overall effectiveness as a monotherapy to treat obesity is relatively modest following 4-8 week treatments in DIO mice (≈4.9%) [29], rats (≈8.7%) [29] and rhesus monkeys (≈3.3%) [19] thus making it more suited as a combination therapy with other drugs that work through other mechanisms. Head and colleagues recently reported that systemic OT and the opioid antagonist, naltrexone, resulted in an enhanced reduction of high-fat, high-sugar meal in rats [30]. Recently, we found that hindbrain (fourth ventricle; 4V) OT treatment in combination with systemic treatment with CL 316243, a drug that directly targets beta-3 adrenergic receptors (β3-AR) to increase brown adipose tissue (BAT) thermogenesis [29; 31; 32; 33; 34], resulted in greater weight loss (15.5 ± 1.2% weight loss) than either OT (7.8 ± 1.3% weight loss) or CL 316243 (9.1 ± 2.1% weight loss) alone [35].

The goal of the current study aimed to determine whether systemic OT treatment could be used as an adjunct with the β3-AR agonist, CL 316243, to increase BAT thermogenesis and elicit weight loss in DIO rats when using a more translational route of administration for OT delivery. We hypothesized that systemic OT and β3-AR agonist (CL 316243) treatment would produce an additive effect to reduce body weight and adiposity in DIO rats by decreasing food intake and stimulating BAT thermogenesis. To test this, we determined the effects of systemic (subcutaneous) infusions of OT (50 nmol/day) or vehicle (VEH) when combined with daily systemic (intraperitoneal (IP)) injections of CL 316243 (0.5 mg/kg) or VEH on body weight, adiposity, food intake, brown adipose tissue temperature (T_IBAT_) and thermogenic gene expression.

## Methods

### Animals

Adult male Long-Evans rats [∼ 8-9 weeks old, 292-349 grams at start of high fat dietary (HFD) intervention/∼ 8-10 months old, 526-929 g body weight at study onset] were initially obtained from Envigo (Indianapolis, IN) and maintained for at least 4 months on a high fat diet (HFD) prior to study onset. All animals were housed individually in Plexiglas cages in a temperature-controlled room (22±2°C) under a 12:12-h light-dark cycle. All rats were maintained on a 1 a.m./1 p.m. light cycle (lights on at 1 a.m./lights off at 1 p.m.). Rats had *ad libitum* access to water and a HFD providing 60% kcal from fat (approximately 6.8% kcal from sucrose and 8.9% of the diet from sucrose) (D12492; Research Diets, Inc., New Brunswick, NJ). The research protocols were approved both by the Institutional Animal Care and Use Committee of the Veterans Affairs Puget Sound Health Care System (VAPSHCS) and the University of Washington in accordance with NIH’s *Guide for the Care and Use of Laboratory Animals* (NAS, 2011) [36]. We have used the ARRIVE Essential 10 checklist for reporting animal studies.

### Drug Preparation

Fresh solutions of OT acetate salt (Bachem Americas, Inc., Torrance, CA) were solubilized in sterile water. Each minipump was placed into a test tube containing sterile 0.9% saline and then into a water bath at 37° C for approximately 40 hours prior to minipump implantation based on manufacturer’s recommended priming instructions for ALZET® model 2004 minipumps. CL 316243 (Tocris/Bio-Techne Corporation, Minneapolis, MN) was solubilized in sterile water each day of each experiment. CL 316243 was used in place of the FDA-approved β3-AR agonist, Mirabegron, due to the well-established effects of CL 316243 on lipolysis in isolated white adipocytes cells [37] and on BAT thermogenesis and energy expenditure in rodent models in our lab [35] and other labs [33; 34; 38; 39]. In contrast to CL 316243, Mirabegron is not water soluble.

### Subcutaneous implantations of osmotic minipumps

The approach for implanting minipumps has been described previously [35]. Briefly, animals received a subcutaneous implantation of osmotic minipump (model 2004, DURECT Corporation Cupertino, CA) one week prior to CL 316243 treatment as previously described [35]. Animals were treated once with the analgesic ketoprofen (2 mg/kg; Fort Dodge Animal Health) at the completion of the minipump implantations.

### Implantation of implantable telemetry devices (PDT-4000 HR E-Mitter or PDT-4000) into the abdominal cavity

Animals were anesthetized with isoflurane and subsequently received implantations of a sterile PDT-4000 HR E-Mitter (26 mm long × 8 mm wide; Starr Life Sciences Company) or PDT-4000 E-Mitter (23 mm long × 8 mm wide) into the intraperitoneal cavity. The abdominal opening was closed with 4-0 Vicryl absorbable suture and the skin was closed with 4-0 monofilament nonabsorbable suture. Vetbond glue was used to seal the wound and bind any tissue together between the sutures. Animals were treated with the analgesic ketoprofen (2 mg/kg; Fort Dodge Animal Health; once/day for 3 consecutive days, including day of surgery) and the antibiotic enrofloxacin (5 mg/kg; Bayer Healthcare LLC., Animal Health Division Shawnee Mission, KS, United States; once per day for 4 consecutive days, including day of surgery) at the completion of the intra-abdominal implantations. Sutures were removed within two weeks after the PDT-4000 HR E-Mitter implantation. All PDT-4000 HR and PDT-4000 E-Mitters were confirmed to have remained within the abdominal cavity at the conclusion of the study.

### Implantation of temperature transponders underneath interscapular brown adipose tissue (IBAT)

The approach for implanting temperature transponders has been described previously [35]. Rats were anesthetized with isoflurane prior to having the dorsal surface along the upper midline of the back shaved and scrubbed with 70% ethanol followed by betadine swabs as previously described [29]. Following an incision (1 “) along the midline of the interscapular area a temperature transponder (14 mm long/2 mm wide) (HTEC IPTT-300; Bio Medic Data Systems, Inc., Seaford, DE) was implanted underneath the left IBAT pad as previously described [29; 40; 41]. The transponder was subsequently secured in place by suturing it to the brown fat pad with sterile silk suture. The interscapular incision was closed with Nylon sutures (5-0), which were removed in awake animals approximately 10-14 days post-surgery. Animals were treated once with the analgesic ketoprofen (2 mg/kg; Fort Dodge Animal Health) at the completion of the temperature transponder implantations. HTEC IPTT-300 transponders were used in place of IPTT-300 transponders to enhance accuracy in our measurements as previously described [29].

### Acute IP injections and measurements of T_IBAT_

CL 316243 (or saline vehicle; 0.1 ml/kg injection volume) was administered immediately prior to the start of the dark cycle following 4 hours of food deprivation. Animals remained without access to food for an additional 1 (**Study 3**) or 4 h (**Studies 1-2**) during the course of the T_IBAT_ measurements. The purpose of the fast was to minimize the potential confound of diet-induced thermogenesis [42] on T_IBAT_, which was a key measurement in our studies. A handheld reader (DAS-8007-IUS Reader System; Bio Medic Data Systems, Inc.) was used to collect measurements of T_IBAT_. Measurements were taken under dimmed red light.

### Body Composition

Determinations of lean body mass and fat mass were made on un-anesthetized rats by quantitative magnetic resonance using an EchoMRI 4-in-1-700^TM^ instrument (Echo Medical Systems, Houston, TX) at the VAPSHCS Rodent Metabolic Phenotyping Core.

Measurements were taken prior to subcutaneous minipump implantations as well as at the end of the infusion period.

## Study Protocols

### Study 1: Determine the dose-response effects of systemic CL 316243 on body weight, energy Intake, T_IBAT_, core temperature and gross motor activity in male DIO rats

Rats (N=10 at study onset) (∼ 9 mo old; 517-823 g at start of study) were fed *ad libitum* and maintained on HFD for approximately 8 months prior to being implanted with temperature transponders (HTEC IPTT-300) underneath the left IBAT depot. Following a 2-week period, animals were subsequently implanted with a PDT-4000 HR E-Mitter telemetry device into the abdominal cavity. Following a 3-week recovery period, CL 316243 or vehicle was administered once per animal at approximately 1-week intervals so that each animal served as its own control. T_IBAT_ was measured daily at baseline (-4 h; 9:00 a.m.), immediately prior to IP injections (0 h; 12:45-1:00 p.m.), and at 0.25, 0.5, 0.75, 1, 20 and 24-h post-injection. The doses of CL 316243 (0.01, 0.1, 0.5 and 1 mg/kg) were selected based on doses of CL 316243 found to be effective at increasing T_IBAT_ when administered intraperitoneally [35] into DIO rats.

*Changes of Core Temperature and Gross Motor Activity.* The protocol for measuring core temperature and gross motor activity has been described previously [43]. Briefly, core temperature (surrogate marker of energy expenditure) and gross motor activity were recorded non-invasively in unanesthetized rats in the home cage every 30 sec throughout the study.

### Study 2: Determine the dose-response effects of systemic infusion of OT (16 and 50 nmol/day) on body weight, adiposity, energy intake and kaolin intake in male DIO rats

Rats (N=20 at study onset) (∼ 10.5 mo old; 577-962 g at start of study) were fed *ad libitum* and maintained on HFD for approximately 8.5 months prior to prior to being implanted with a temperature transponder underneath the left IBAT depot. Following a 2-week period, animals were matched for both body weight and adiposity prior to being implanted with minipumps. Rats were subsequently maintained on a daily 4-h fast and received minipumps to infuse vehicle or OT (16 or 50 nmol/day) over 29 days. These doses were selected based on a dose of OT found to be effective at reducing body weight or body weight gain when administered subcutaneously [19; 20] or into the 4V [44] of HFD-fed rats. Daily food intake and body weight were collected across the 29-day infusion period.

### Study 3: Effect of chronic systemic OT infusions (50 nmol/day) and systemic beta-3 receptor agonist (CL 316243) administration (0.5 mg/kg) on body weight, body adiposity, energy intake and T_IBAT_ in male DIO rats

Rats (N=43 at study onset) (∼ 10 mo old; 526-929 g at start of study) were fed *ad libitum* and maintained on HFD for approximately 9 months prior to receiving implantations of temperature transponders underneath the left IBAT pad. Following a 1-week recovery period, a subset of animals (N=15) was implanted with a PDT-4000 E-Mitter telemetry device into the abdominal cavity or received sham implantations (N=10). Following up to a 1-month recovery period, animals were matched for both body weight and adiposity and were subsequently implanted with minipumps to infuse vehicle or OT (50 nmol/day) over 29 days, respectively. After having matched animals for OT-elicited reductions in body weight (infusion day 7), DIO rats subsequently received single daily IP injections of VEH or CL 316243 (0.5 mg/kg). We selected this dose of CL 316243 because it reduced energy intake and body weight gain and elevated both T_IBAT_ and core temperature and (**Study 1**). Importantly, we found that this dose of CL 3162243 and CNS administration of OT were found to produce an additive effect on weight loss in DIO rats [35]. In addition, we selected the dose of OT (50 nmol/day) because it produced comparable weight loss to that of OT alone in **Study 1**. T_IBAT_ was measured daily at baseline (-4 h; 9:00 a.m.), immediately prior to IP injections (0 h; 12:45-1:00 p.m.), and at 0.25, 0.5, 0.75, 1, 20 and 24-h post-injection. In addition, daily food intake and body weight were collected across the 29-day infusion period. Data from animals that received the single dose of CL-316243 were analyzed over the 29-day infusion period.

### Kaolin measurements

Kaolin (K50001, Research Diets, Inc.) intake (g) was assessed over 29 days following implantation of minipumps containing vehicle (saline) or OT (16 or 50 nmol/day). Placement of kaolin and high fat diet was reversed every other day within each treatment condition.

### Adipose tissue processing for adipocyte size and UCP-1 analysis

Inguinal white adipose tissue (IWAT) and epididymal white adipose tissue (EWAT) depots were collected at the end of the infusion period in rats from **Study 3**. Rats from each group were euthanized following a 3-h fast. Rats were euthanized with intraperitoneal injections of ketamine cocktail [ketamine hydrochloride (214.3 mg/kg), xylazine (10.71 mg/kg) and acepromazine (3.3 mg/kg) in an injection volume up to 2 mL/rat] and transcardially exsanguinated with PBS followed by perfusion with 4% paraformaldehyde in 0.1 M PBS. Adipose tissue (IBAT, IWAT, and EWAT) was dissected and placed in 4% paraformaldehyde-PBS for 24 h and then placed in 70% ethanol (EtOH) prior to paraffin embedding. Sections (5 μm) sampled were obtained using a rotary microtome, slide-mounted using a floatation water bath (37°C), and baked for 30 min at 60°C to give approximately 15-16 slides/fat depot with two sections/slide.

### Adipocyte size analysis and immunohistochemical staining of UCP-1

Adipocyte size analysis was performed on deparaffinized, airdried, unstained and uncovered sections. Slightly underexposed photographs of dry sectioned produced clear, highly contrasted black and white images suitable for a built-in particle counting method of ImageJ software (National Institutes of Health, Bethesda, MD). Images were first converted to 16-bit files and then modified and analyzed with a cell shape factor 0.35-1 (a shape factor of 0 represents a straight line and a shape factor of 1 indicates a circle) (methods modified from [45]). Fixed (4% PFA), paraffin-embedded adipose tissue was sectioned and stained with a-primary rabbit anti-UCP-1 antibody (1:100; Abcam, Cambridge, MA (#ab10983/RRID: AB_2241462)] as has been previously described in lean C57BL/6J mice [46] and both lean and DIO C57BL/6 mice after having been screened in both IBAT and IWAT of Ucp1^+/-^ and Ucp1^-/-^ mice [47]. Immunostaining specificity controls included omission of the primary antibody and replacement of the primary antibody with normal rabbit serum at the same dilution as the respective primary antibody. Area quantification for UCP-1 staining was performed on digital images of immunostained tissue sections using image analysis software (Image Pro Plus software, Media Cybernetics, Rockville, MD, USA). Slides were visualized using bright field on an Olympus BX51 microscope (Olympus Corporation of the Americas; Center Valley, PA) and photographed using a Canon EOS 5D SR DSLR (Canon U.S.A., Inc., Melville, NY) camera at 100X magnification. Values for each tissue within a treatment were averaged to obtain the mean of the treatment group.

### Blood collection

Blood was collected from 4-h (**Study 1**) or 6-h fasted rats (**Studies 2-3**) within a 2-h window towards the end of the light cycle (10:00 a.m.-12:00 p.m.) as previously described in DIO CD^®^ IGS rats and mice [25; 29]. Animals from **Study 3** were euthanized at 2-h post-CL 316243 or VEH treatment. Treatment groups were counterbalanced at time of euthanasia to avoid time of day bias. Blood samples [up to 3 mL] were collected immediately prior to transcardial perfusion by cardiac puncture in chilled K2 EDTA Microtainer Tubes (Becton-Dickinson, Franklin Lakes, NJ). Whole blood was centrifuged at 6,000 rpm for 1.5-min at 4°C; plasma was removed, aliquoted and stored at −80°C for subsequent analysis.

### Plasma hormone measurements

Plasma leptin and insulin were measured using electrochemiluminescence detection [Meso Scale Discovery (MSD^®^), Rockville, MD] using established procedures [29; 48]. Intra-assay coefficient of variation (CV) for leptin was 2.7% and 3.2% for insulin. The range of detectability for the leptin assay is 0.137-100 ng/mL and 0.069-50 ng/mL for insulin. Plasma fibroblast growth factor-21 (FGF-21) (R&D Systems, Minneapolis, MN) and irisin (AdipoGen, San Diego, CA) levels were determined by ELISA. The intra-assay CV for FGF-21 and irisin were 4.5% and 8.4%, respectively; the ranges of detectability were 31.3-2000 pg/mL (FGF-21) and 0.078-5 μg/mL (irisin). Plasma adiponectin was also measured using electrochemiluminescence detection Meso Scale Discovery (MSD^®^), Rockville, MD] using established procedures [29; 48]. Intra-assay CV for adiponectin was 1.1%. The range of detectability for the adiponectin assay is 2.8-178 ng/mL. The data were normalized to historical values using a pooled plasma quality control sample that was assayed in each plate.

### Blood glucose and lipid measurements

Blood was collected for glucose measurements by tail vein nick following a 4 (**Study 1**) or 6-h fast (**Studies 2-3**) and measured with a glucometer using the AlphaTRAK 2 blood glucose monitoring system (Abbott Laboratories, Abbott Park, IL) [49]. Tail vein glucose was measured at 2-h post-CL 316243 or VEH treatment (**Study 3**). Total cholesterol (TC) [Fisher Diagnostics (Middletown, VA)] and free fatty acids (FFAs) [Wako Chemicals USA, Inc., Richmond, VA)] were measured using an enzymatic-based kits. Intra-assay CVs for TC and FFAs were 1.4 and 2.3%, respectively. These assay procedures have been validated for rodents [50].

### Tissue collection for quantitative real-time PCR (qPCR)

IBAT, EWAT and IWAT tissue was collected from 4 (**Study 1**) or 6-h fasted rats (**Study 3**). In addition, animals from **Study 3** were euthanized at 2-h post-CL 316243 (0.5 mg/kg) or VEH administration. IBAT, EWAT and IWAT were collected within a 2-h window towards the end of the light cycle (10:00 a.m.-12:00 p.m.) as previously described in DIO CD^®^ IGS/Long-Evans rats and C57BL/6J mice [25; 29; 44]. Tissue was rapidly removed, wrapped in foil and frozen in liquid N2. Samples were stored frozen at-80°C until analysis.

### qPCR

RNA extracted from samples of IBAT, EWAT and IWAT (**Study 3**) were analyzed using the RNeasy Lipid Mini Kit (Qiagen Sciences Inc, Germantown, MD) followed by reverse transcription into cDNA using a high-capacity cDNA archive kit (Applied Biosystems, Foster City, CA). Quantitative analysis for relative levels of mRNA in the RNA extracts was measured in duplicate by qPCR on an Applied Biosystems 7500 Real-Time PCR system (Thermo Fisher Scientific, Waltham, MA). The TaqMan® probe for rat *Nono* (Rn01418995_g1), *Ucp1 (*catalog no. Rn00562126_m1), *Adrb1 (*catalog no. Rn00824536_s1), *Adrb3 (*catalog no. Rn01478698_g1), type 2 deiodinase (*Dio2)* (catalog no. Rn00581867_m1), G-protein coupled receptor 120 (*Gpr120*; catalog no. Rn01759772_m1), cell death-inducing DNA fragmentation factor α-like effector A (*Cidea*; catalog no. Rn04181355_m1), peroxisome proliferator-activated receptor gamma coactivator 1α (*Ppargc1a*; catalog no. Rn00580241_m1) and PR domain containing 16 (*Prdm16*; catalog no. Rn01516224_m1) were acquired from Thermo Fisher Scientific (Thermo Fisher Scientific Gene Expression Assay probes). Relative amounts of target mRNA were determined using the Comparative C_T_ or 2-^ΔΔC^ method [51] following adjustment for the housekeeping gene, *Nono*. Specific mRNA levels of all genes of interest were normalized to the cycle threshold value of *Nono* mRNA in each sample and expressed as changes normalized to controls (vehicle/vehicle treatment).

### Verification of transponder and telemetry device (E-Mitter) placement

All temperature transponders, PDT-4000 HR E-Mitters and PDT-4000 E-Mitters were confirmed to have remained underneath the IBAT depot and within the abdominal cavity, respectively, at the conclusion of the study.

## Statistical Analyses

All results are expressed as means ± SE. Comparisons between multiple groups involving between-subjects designs were made using one-or two-way ANOVA as appropriate, followed by a post-hoc Fisher’s least significant difference test. Comparisons involving within-subjects designs were made using a one-way repeated-measures ANOVA followed by a post-hoc Fisher’s least significant difference test. Analyses were performed using the statistical program SYSTAT (Systat Software, Point Richmond, CA). Differences were considered significant at *P*<0.05, 2-tailed.

## Results

### Study 1: Determine the dose-response effects of acute systemic CL 316243 on body weight, energy intake, T_IBAT_, core temperature and gross motor activity in male DIO rats

The goal of this study was to identify a dose range of the beta-3 receptor agonist, CL 316243, that resulted in weight loss, reduced energy intake and an elevation in IBAT and core temperature in DIO rats. The effective dosing data from this study was used to select a dose range (0.01-1 mg/kg, IP) for use in the subsequent chronic dose escalation study (**Study 3**). By design, DIO rats were obese as determined by both body weight (782.1±22.2 g) and adiposity (288.4±12.4 g fat mass; 36.8±0.8% adiposity) after maintenance on the HFD for approximately 8 months.

### Body weight

As in our previous study in DIO rats [35], there was an overall significant main effect of CL 316243 to reduce body weight gain at 20-[(F(4,36) = 4.347, *P*=0.006], 44-[(F(4,36) = 4.734, *P*=0.004], 68-[(F(4,36) = 7.316, *P*<0.01], 92-[(F(4,36) = 3.858, *P*=0.010], 116-[(F(4,36) = 8.203, *P*<0.01], 140-[(F(4,36) = 7.531, *P*<0.01], 164-[(F(4,36) = 6.100, *P*=0.001] and 188-h post-injection [(F(4,36) = 5.129, *P*=0.002].

Specifically, the highest dose (1 mg/kg) reduced body weight gain across all post-injection time intervals (*P*<0.05 vs VEH) (**Figure 1A**). The second highest dose of CL 316243 (0.5 mg/kg) also reduced weight gain at 20, 44, 68, 92, 1116, 140, and 164-h post-injection. A lower dose of CL 31243 (0.1 mg/kg) was also effective at reducing weight gain at 20-, 44-, 68-, 116, 140, 164 and 188-h post-0injection. The lowest dose (0.01 mg/kg) reduced body weight gain at 44-, 116-h post-injection and tended to reduce body weight gain at 68-h post-injection.

**Figure 1.**
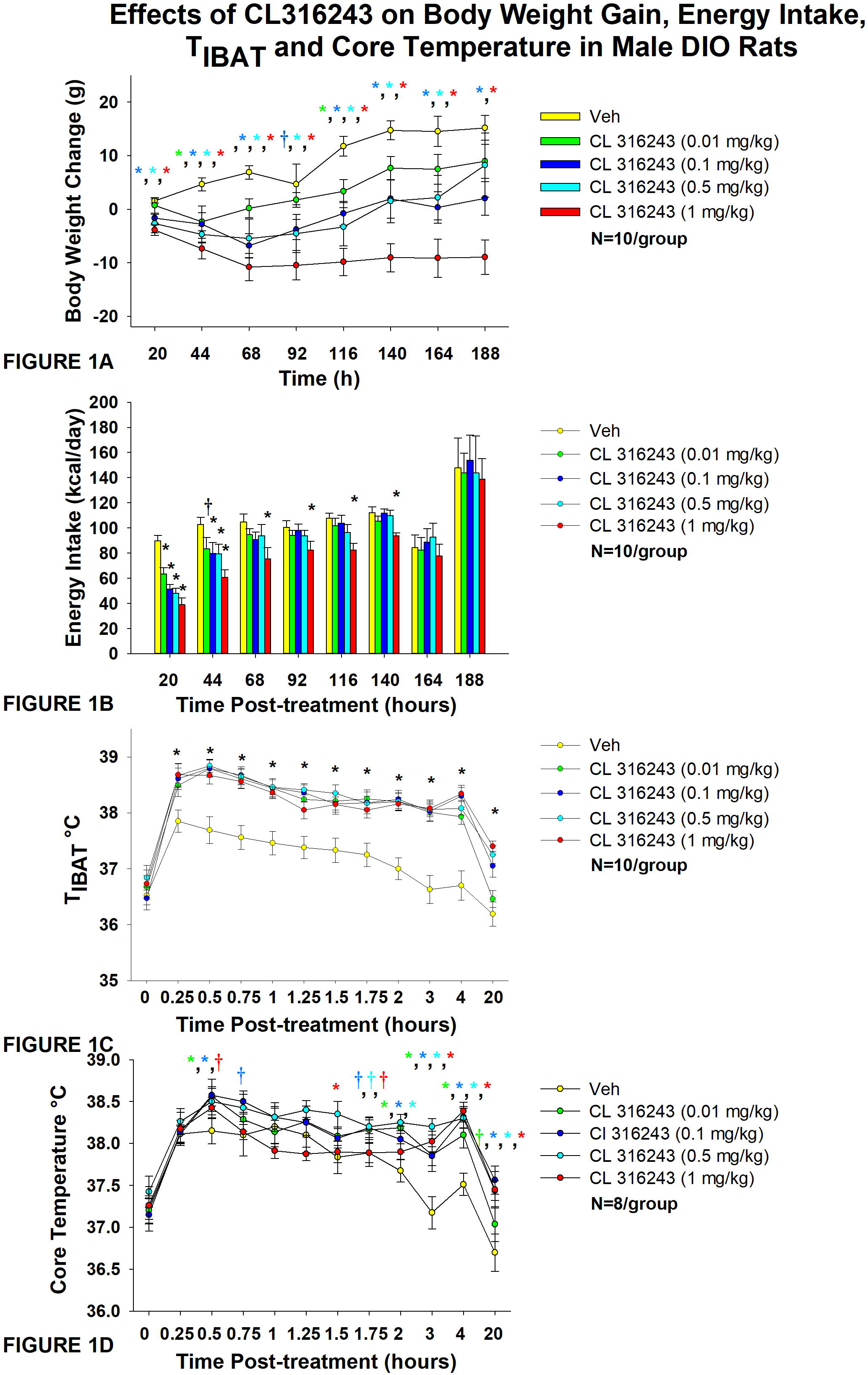
Dose-response effects of the beta-3 receptor agonist, CL 316243, on body weight, energy intake, T_IBAT_, core temperature and gross motor activity in male DIO rats. Ad libitum fed rats were either maintained on HFD (60% kcal from fat; N=10-12/group) for approximately 8 months prior to receiving IP injections of CL 316243 (0.01-1 mg/kg) or vehicle (sterile water) where each animal received each treatment approximately once per week. *A*, Effect of CL 316243 on body weight change in DIO rats *B*, Effect of CL 316243 on energy intake (kcal/day) in DIO rats *C*, Effect of CL 316243 on T_IBAT_ in DIO rats; *D*, Effect of CL 316243 on core temperature in DIO rats. Data are expressed as mean ± SEM. \**P*<0.05, †0.05<*P*<0.1 CL 316243 vs. vehicle.

### Energy intake

As in our previous study in DIO rats [35], there was a significant main effect of CL 316243 to reduce energy intake at 20-h [(F(4,36) = 17.431, *P*<0.01], 44-h [(F(4,36) = 3.688, *P*=0.013], 116-h [(F(4,36) = 3.094, *P*=0.027] and 140-h post-injection [(F(4,36) = 4.416, *P*=0.007]. In addition, there was a near significant effect at 68-h post-injection [(F(4,36) = 2.131, *P*=0.097].

Specifically, CL 316243 reduced 20-h energy intake at 0.01, 0.1, 0.5 and 1 mg/kg (*P*<0.05) by 28.4 ± 5.5, 41.4 ± 5.4, 45.0 ± 6.0, and 56.5 ± 5.8% (**Figure 1B**). The higher doses (0.1, 0.5 and 1 mg/kg) also reduced energy intake by 18.0 ± 11.1, 17.8 ± 12.1, and 37.8 ± 7.7% (*P*<0.05) at 44-h post-injection (*P*<0.05). Furthermore, the highest dose (1 mg/kg) reduced energy intake by 25.7 ± 10.3, 13.0 ± 11.6, 22.8 ± 5.7 and 13.0 ± 11.6% at 68-, 92-, 116-and 140-h post-injection (*P*<0.05).

### T_IBAT_

As in our previous study in DIO rats [35], there was a significant main effect of CL 316243 to elevate T_IBAT_ at 0.25-[(F(4,36) = 22.639, *P*<0.01], 0.5-[(F(4,36) = 31.681, *P*<0.01], 0.75-[(F(4,36) = 19.777, *P*<0.01], 1-[(F(4,36) = 14.901, *P*<0.01], 1.25-[(F(4,36) = 19.606, *P*<0.01], 1.5-[(F(4,36) = 12.651, *P*<0.01], 1.75-[(F(4,36) = 18.074, *P*<0.01], 2-[(F(4,36) = 25.583, *P*<0.01], 3-[(F(4,36) = 23.501, *P*<0.01], 4-[(F(4,36) = 30.853, *P*<0.01] and 16-h post-injection [(F(4,36) = 13.296, *P*<0.01]. We also found a significant effect of time [(F(10,450) = 99.883, *P*<0.01] and a significant interactive effect between time and dose [(F(40,450) = 2.635, *P*<0.01] across 11 time points over the 16-h measurement period.

Specifically, systemic injections of CL 316243 (0.01-1 mg/kg) (**Figure 1C**) increased T_IBAT_ at all time points between 0.25-h and 4-h post-injection. The higher doses (0.1-1 mg/kg) also increased T_IBAT_ at 16-h post-injection (*P*<0.05).

### Core temperature

There was a significant main effect of CL 316243 to elevate core temperature at 2-[(F(4,28) = 5.833, P<0.01], 3-[(F(4,28) = 7.394, *P*<0.01], 4-[(F(4,28) = 9.207, *P*<0.01], 5-[(F(4,28) = 3.534, *P*=0.019], 14-[(F(4,28) = 4.497, *P*=0.006], 16-[(F(4,28) = 7.136, *P*<0.01], 22-[(F(4,28) = 3.382, *P*=0.022], 24-[(F(4,28) = 14.967, P<0.01] and 40-h post-injection [(F(4,28) = 4.718, *P*=0.005].

CL 316243 produced a near significant main effect to elevate core temperature at 1.75-[(F(4,28) = 2.211, *P*=0.093] and 6-h post-injection [(F(4,28) = 2.419, *P*=0.072]. We also found a significant effect of time [(F(12,420) = 25.873, *P*<0.01] and a significant interactive effect between time and dose [(F(48,420) = 1.830, *P*=0.001] across 13 time points over the initial 16-h measurement period.

Specifically, systemic injections of CL 316243 increased core temperature at 0.5- (0.01 and 0.1 mg), 1.5- (0.5 mg/kg), 2-(0.01, 0.1 and 0.5 mg/kg), 3- (0.01, 0.1, 0.5 and 1 mg/kg), 4- (0.01, 0.1, 0.5 and 1 mg/kg), 5- (0.1 and 1 mg/kg), 6- (0.1, 0.5 and 1 mg/kg), 14- (0.01, 0.1, 0.5 and 1 mg/kg), 16- (0.1, 0.5 and 1 mg/kg), 22- (0.5 and 1 mg/kg), 24-(0.5 and 1 mg/kg), and 40-h post-injection (1 mg/kg).

There was also a near significant effect of CL 316243 to stimulate core temperature at 0.5-h (0.5 mg/kg), 0.75-h (0.1 mg/kg), 1.75-h (0.01, 0.1 and 0.5 mg/kg), 5-h (0.01 and 0.5 mg/kg), 6-h (0.01 mg/kg), 12-h (0.1 and 0.5 mg/kg), 16-(0.01 mg/kg), 20-h post-injection (0.5 mg/kg) and 24-h post-injection (0.01 mg/kg).

Two animals were removed from the core temperature and gross motor activity analysis due to defective telemetry devices.

### Gross motor activity

There was a significant main effect of CL 316243 to reduce gross motor activity at 1-h post-injection [(F(4,28) = 5.603, *P*=0.002]. There was also a near significant main effect of CL 316243 to reduce gross motor activity at 0.75-h post-injection [(F(4,28) = 2.375, *P*=0.076].

Specifically, CL 316243 reduced gross motor activity at 0.75-(0.01, 0.1 and 0.5 mg/kg), and 1-h post-injection (0.01, 0.1, 0.5 and 1 mg/kg) (*P*<0.05; data not shown). There was also a near significant effect of CL 316243 (0.01 and 1 mg/kg) to reduce core temperature at 22-h post-injection (0.05<*P*<0.01). Otherwise, CL 316243 was ineffective at altering gross motor activity at any other time point (*P*=NS vs vehicle; data not shown).

### Study 2: Determine the dose-response effects of systemic infusion of OT (16 and 50 nmol/day) on body weight, adiposity, energy intake and kaolin intake in male DIO rats

By design, rats were diet-induced obese (DIO) as determined by both body weight (771 ±24.5 g) and adiposity (312.3±16.7 g fat mass; 40.1±1.1% adiposity) after maintenance on the HFD for approximately 8.5 months. Following temperature transponder implantations and prior to minipump implantations, groups were again matched for body weight and adiposity (vehicle: 760±56.5 grams/36.8±1.7% fat/283.3±31.9 g fat mass); OT (16 nmol/day): 758.6±39.9 grams/36.8±2.0% fat/282.0±28.1 g fat mass; OT (50 nmol/day): 763.6±45.7 grams/36.6±1.8% fat/284.5±31.1 g fat mass. There was no difference in body weight [(F(2,17) = 0.027, *P*=NS)] or percent adiposity [(F(2,17) = 0.003, *P*=NS)] between groups prior to treatment onset. As expected, body weight of DIO rats remained stable over the month of vehicle treatment relative to pre-treatment [(F(1,5) = 2.865, *P*=0.151)] (**Figure 2A**). In contrast to vehicle treatment, systemic OT (16 nmol/day) resulted in a significant reduction of body weight relative to OT pre-treatment [(F(1,6) = 140.799, *P*<0.01)] (**Figure 2A**; *P*<0.05). Furthermore, SC OT, at a 3-fold higher dose (50 nmol/day), also resulted in a significant reduction of body weight relative to pre-treatment [(F(1,6) = 47.271, *P*<0.01)].

**Figure 2A-D:**
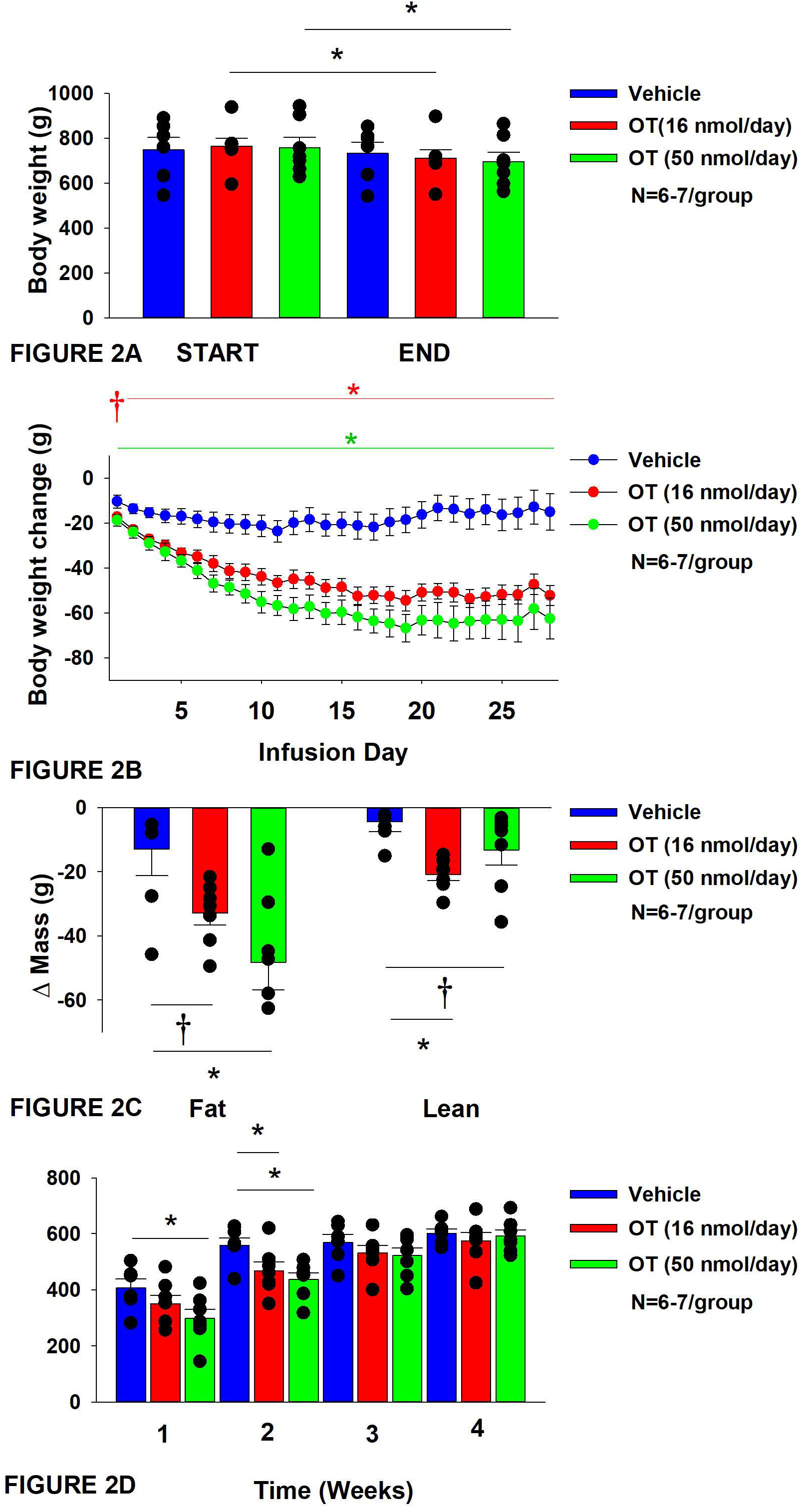
Determine the dose-response effects of systemic infusions of OT (16 and 50 nmol/day) on body weight, adiposity and energy intake in DIO rats. *A*, Rats were maintained on HFD (60% kcal from fat; N=6-7/group) for approximately 5.5 months prior to being implanted with temperature transponders and allowed to recover for 1-2 weeks prior to being implanted with subcutaneous minipumps. *A*, Effect of chronic subcutaneous OT or vehicle on body weight in DIO rats; *B*, Effect of chronic subcutaneous OT or vehicle on body weight change in DIO rats; *C*, Effect of chronic subcutaneous OT or vehicle on change in fat mass and lean mass in DIO rats; *D*, Effect of chronic subcutaneous OT or vehicle on change in weekly energy intake (kcal/week) in DIO rats. Data are expressed as mean ± SEM. \**P*<0.05, †0.05<*P*<0.1 OT vs. vehicle.

In addition, SC OT (16 and 50 nmol/day) was able to reduce weight gain (**Figure 2B**) relative to vehicle treatment throughout the 28-day infusion period. SC OT (50 nmol/day), at a dose that was at least 3-fold higher than the centrally effective dose (16 nmol/day), reduced weight gain throughout the entire 28-day infusion period. SC OT (16 nmol/day) treated rats had reduced weight gain between days 2-29 (*P*<0.05) while SC OT (50 nmol/day) reduced weight gain between days 1-29 (*P*<0.05). There was an overall effect of OT to reduce relative fat mass (pre-vs post-intervention) [(F(2,17) = 6.052, *P*=0.010)]. SC OT (50 nmol/day) reduced fat mass (*P*<0.05) and there was also a tendency for the lower dose (16 nmol/day) to reduce relative fat mass (*P*=0.066) (**Figure 2C**; *P*<0.05).

There was also an overall effect of OT to reduce relative lean mass (pre-vs post-intervention) [(F(2,17) = 5.572, *P*=0.014)]. Specifically, SC OT (16 nmol/day) reduced relative lean mass at the lower dose (16 nmol/day; *P*<0.01) while the higher dose (50 nmol/day) tended to reduce relative lean mass (*P*=0.090). Note that there was no significant reduction in total fat mass or lean mass (*P*=NS).

The changes in body weight and relative fat mass were not associated with any changes in plasma leptin, insulin, glucose or total cholesterol (**Table 1**). These effects that were mediated, at least in part, by a modest reduction of energy intake that was apparent during weeks 1 (50 nmol/day) and 2 (16 and 50 nmol/day) of OT treatment (**Figure 2D**; *P*<0.05). The reduction of energy intake does not appear to be due to an aversive effect of systemic OT (16 or 50 nmol/day), since there was no effect on kaolin consumption relative to vehicle-treated DIO rats (*P*=NS; data not shown).

**Table 1.** Plasma measurements following systemic infusions of OT (16 and 50 nmol/day) or vehicle DIO rats. Data are expressed as mean ± SEM. \**P*<0.05 OT vs. vehicle (N=6-7/group).

| **SC** | **VEH** | **OT (16 nmol/day)** | **OT (50 nmol/day)** |
| --- | --- | --- | --- |
| **Leptin (ng/mL)** | 65.9 ± 7.8 | 71.3 ± 8.2 | 59.6 ± 7.7 |
| **Insulin (ng/mL)** | 4.7 ± 1.7 | 3.4 ± 0.9 | 4.7 ± 1.3 |
| **Blood Glucose (mg/dL)** | 137.3 ± 4.5 | 160.4 ± 8.3* | 157.7 ± 7.3 |
| **Total Cholesterol (mg/dL)** | 108.1 ± 10.4 | 118.4 ± 13.4 | 105.5 ± 11.5 |

**Blood was collected in 4-h fasted rats by tail vein nick (glucose) or cardiac stick (leptin, insulin and total cholesterol)**
**\*P<0.05 vs vehicle**

Energy intake data from a vehicle-treated animal during week 3 was deleted on account of an error that was made when recording the data. Kaolin data from a subset of animals was excluded on account of these animals shredding the kaolin diet (2 measurements during week 2, 1 measurement during week 3 and 1 measurement during week 4) and being significant outliers (Grubb’s test for outliers).

### Study 3: Effect of chronic systemic OT infusions (50 nmol/day) and systemic β3-AR agonist (CL 316243) administration (0.5 mg/kg) on body weight, body adiposity, energy intake and kaolin intake in male DIO rats

The goal of this study was to determine the effects of chronic OT treatment (single dose identified from **Study 2**) in combination with a single dose of the β3-AR agonist, CL 316243, on body weight and adiposity in DIO rats. By design, DIO rats were obese as determined by both body weight (804±14 g) and adiposity (310±11 g fat mass; 38.3±4.7% adiposity) after maintenance on the HFD for approximately 7 months. Prior to the onset of CL 316243 treatment on infusion day 7, both OT treatment groups were matched for OT-elicited reductions of weight gain.

OT and CL 316243 alone reduced body weight by ≍ 7.8±1.3% (P<0.05) and 9.1±2.3% (*P*<0.05), respectively, but the combined treatment produced more pronounced weight loss (pre-vs post-intervention) (15.5±1.2%; *P*<0.05) (**Figure 3A**) than either treatment alone (*P*<0.05). OT alone tended to reduce weight gain on days 16-28 (0.05<*P*<0.1) while CL 316243 alone tended to reduce or reduced weight gain on day 24 (0.05<*P*<0.1), day 25 (*P*=0.05), and days 26-28 (*P*<0.05) (**Figure 3B**). OT and CL 316243 together tended to reduce weight gain on day 9 (0.05<*P*<0.1) reduced weight gain on days 10-28 (*P*<0.05). The combination treatment appeared to produce a more pronounced reduction of weight gain relative to OT alone on day 25 (0.05<P<0.1) and this reached significance on days 26-28 (*P*<0.05).

**Figure 3A-E:**
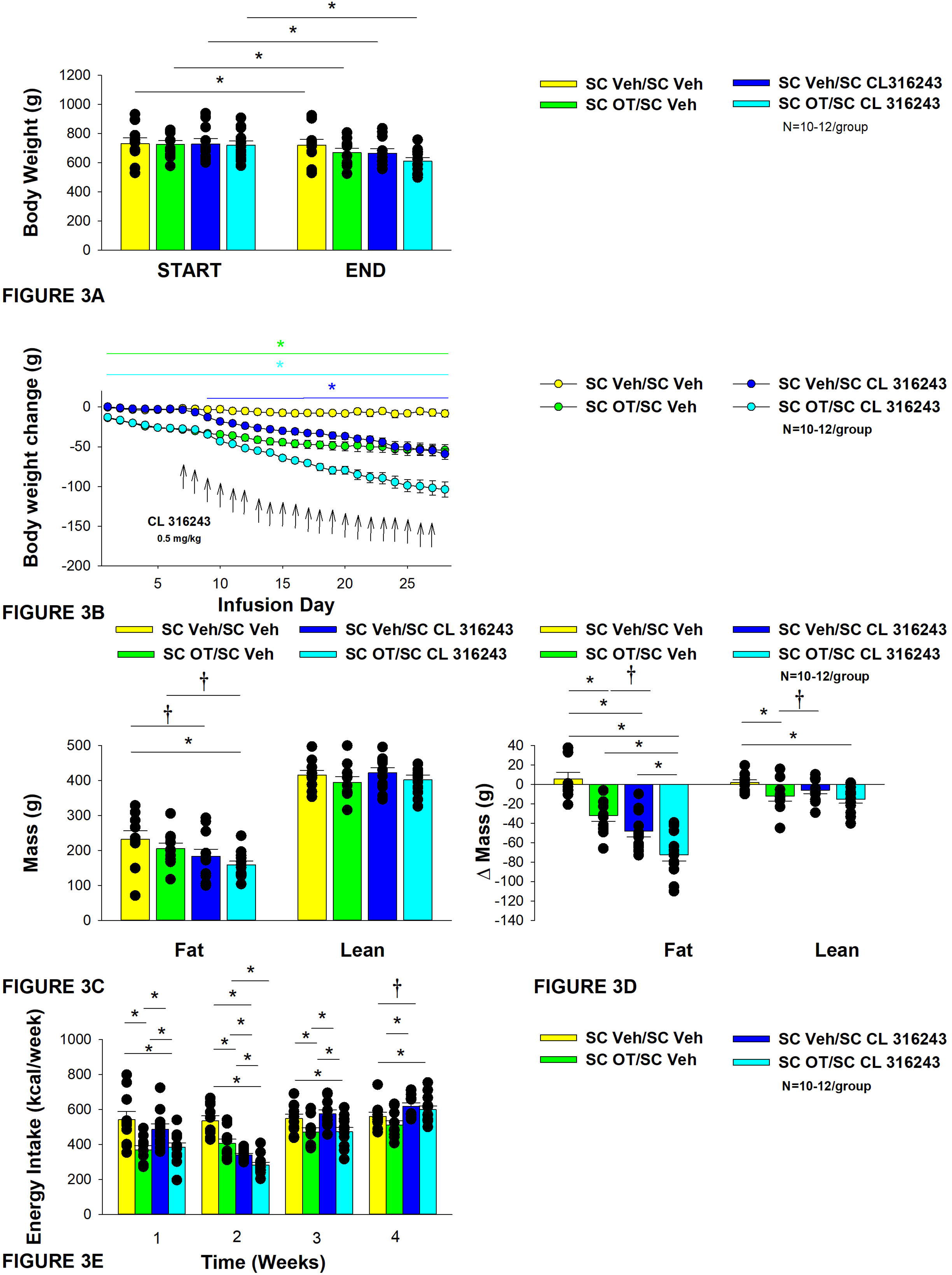
Effect of chronic systemic OT infusions (50 nmol/day) and systemic beta-3 receptor agonist (CL 316243) administration (0.5 mg/kg) on body weight, body adiposity and energy intake in male DIO rats. Ad libitum fed rats were either maintained on HFD (60% kcal from fat; N=8-10/group) for approximately 8 months prior to receiving continuous infusions of vehicle or OT (50 nmol/day) in combination with a single dose of CL 316243 (0.5 mg/kg). *A*, Effect of chronic subcutaneous OT or vehicle in combination with systemic CL 316243 or vehicle on body weight in DIO rats; *B*, Effect of chronic subcutaneous OT or vehicle in combination with systemic CL 316243 or vehicle on body weight change in HFD-fed DIO rats; *C*, Effect of chronic subcutaneous OT or vehicle in combination with systemic CL 316243 or vehicle on fat mass and lean mass in DIO rats; *D*, Effect of chronic subcutaneous OT or vehicle in combination with systemic CL 316243 or vehicle on change in fat mass and lean mass in DIO rats; *E*, Effect of chronic subcutaneous OT or vehicle in combination with systemic CL 316243 or vehicle on change in weekly energy intake (kcal/week) in DIO rats. ↑ indicate 1x daily injections. Colored bars represent specific group comparisons vs vehicle. Data are expressed as mean ± SEM. \**P*<0.05, †0.05<*P*<0.1 vs. vehicle or baseline (pre-treatment; Fig. 3A).

In addition, the combination treatment tended to produce a greater reduction of weight gain relative to CL 316243 alone on days 21-22 and 26-28 (0.05<*P*<0.1). While OT alone did not significantly reduce fat mass (*P*=NS), there was a tendency for CL 316243 alone (0.05<*P*<0.1), and the combination of OT and CL 316243 (*P*<0.05) to reduce fat mass without impacting lean body mass (**Figure 3C**; *P*=NS). However, the combination treatment did not result in a significant reduction of fat mass relative to OT alone or CL 316243 alone (*P*=NS). OT and CL 316243 alone did produce a reduction in relative fat mass (pre-vs post-intervention; *P*<0.05). OT also produced a modest reduction in relative lean mass (**Figure 3D**; *P*<0.05). The combination treatment also produced a significant reduction of relative fat mass (*P*<0.05) which exceeded that of OT and CL 316243 alone (*P*<0.05).

Systemic OT treatment alone reduced energy intake during week 1 (*P*<0.05). OT, CL 316243 and the combined treatment were effective at reducing energy intake at week 2 (**Figure 3E**; *P*<0.05). OT and the combined treatment reduced energy intake during week 3 (*P*<0.05) but CL 316243 failed to reduce energy intake during this time (*P*=NS). All treatments were ineffective at reducing energy intake over weeks 3 and 4 (*P*=NS). The reduction of energy intake in response to OT alone, CL 316243 alone or the combined treatment does not appear to be due to an aversive effect, since there was no effect on kaolin consumption relative to vehicle-treated DIO rats (*P*=NS; data not shown).

Two-way ANOVA revealed an overall significant effect of OT [(F(1,39) = 63.434, *P*<0.01)], CL 316243 [(F(1,39) = 74.939, *P*<0.01)] but no significant interactive effect between OT and CL 316243 [(F(1,39) = 0.058, *P*=NS)] on weight loss. Consistent with this finding, two-way ANOVA revealed consistent overall effects of OT and CL-3162343 on reduction of body weight gain between days 10-29 but no significant overall effect. In addition, two-way ANOVA revealed no overall significant effect of OT [(F(1,39) = 2.003, *P*=0.165)], a significant effect of CL 316243 [(F(1,39) = 6.859, *P*=0.012)] and no significant interactive effect between OT and CL 316243 [(F(1,39) = 0.005, *P*=0.943)] on fat mass. There was no significant overall effect of OT [(F(1,39) = 2.069, *P*=0.158)] or CL 316243 [(F(1,39) = 0.217, *P*=0.644)] on lean mass and no interactive effect of OT and CL 316243 [(F(1,39) = 0.004, *P*=0.950)] on lean mass. Lastly, two-way ANOVA revealed an overall significant effect of OT [(F(1,39) = 21.464, *P*<0.01)], CL 316243 [(F(1,39) = 62.681, *P*<0.01)] and a near significant interactive effect between OT and CL 316243 [(F(1,39) = 3.190, *P*=0.082)] on energy intake (week 2).

Overall, these findings suggest an additive effect of OT and CL 316243 to produce sustained weight loss in DIO rats. The effects of the combination treatment on adiposity and energy intake appear to be driven largely by CL 316243 and OT, respectively.

CL 316243 elevated T_IBAT_ on injection day 1 at 0.5, 0.75 and 1-h post-injection (*P*<0.05; **Supplemental Figure 1A**) and tended to elevate T_IBAT_ at 0.25-h post-injection (0.05<*P*<0.1). Similarly, CL 316243, when given in combination with OT, also increased T_IBAT_ at 0.5, 0.75 and 1-h post-injection and tended to elevate T_IBAT_ at 0.25-h post-injection on injection day 1 (0.05<*P*<0.1). Both CL 316243 and CL 316243 + OT treatments elevated T_IBAT_ relative to vehicle treated animals when the T_IBAT_ data from injection day 1 were averaged over 1-h post-injection (*P*<0.05). There was no significant difference in T_IBAT_ response to CL 316243 CL 316243 + OT treatments when the T_IBAT_ data were averaged over 60 min (*P*=NS).

CL 316243 also elevated T_IBAT_ on injection day 22 at 0.25, 0.5, 0.75 and 1-h post-injection (*P*<0.05; **Supplemental Figure 1B**). Similarly, CL 316243, when given in combination with OT, also increased T_IBAT_ at 0.5 and 0.75-h post-injection and tended to elevate T_IBAT_ at 0.25 and 1-h post-injection on injection day 22 (0.05<*P*<0.1). Both CL 316243 and CL 316243 + OT treatments elevated T_IBAT_ relative to vehicle treated animals when the T_IBAT_ data from injection day 22 were averaged over 1-h post-injection (*P*<0.05). There was no significant difference in the T_IBAT_ response to CL 316243 and CL 316243 +OT treatments when the T_IBAT_ data were averaged over 60 min (*P*=NS).

In addition, in a limited number of subjects (N=3-5/group), we found that CL 31243 in combination with OT elevated core temperature at 20-h post-injection relative to vehicle-treated animals (P<0.05; data not shown). CL 316243 alone also tended to elevate 20-h core temperature (P=0.1; data not shown). No changes in 20-h activity occurred in response to the treatments (data not shown).

### Adipocyte size

H&E-stained sections from the four treatment conditions are shown in **Figure 4A-D (EWAT) and Figure 4E-H (IWAT)**. CL 316243 alone (*P*<0.05) and in combination with OT (*P*<0.05) reduced EWAT adipocyte size in DIO rats relative to vehicle treatment (**Supplemental Figure 2A; Figure 5A**). There were no significant differences in the ability of the combined treatment to reduce EWAT adipocyte size relative to CL 316243 alone (*P*=0.156). Similarly, OT and CL 316243 given in combination reduced IWAT adipocyte size whereas there was no significant effect of OT or CL 316343 on adipocyte size when given alone (*P*=NS) (**Supplemental Figure 2B; Figure 5B**). A subset of EWAT (N=6) and IWAT (N=5) samples were excluded from the adipocyte size analysis due to poor tissue quality or an insufficient amount of tissue.

**Figure 4A-D:**
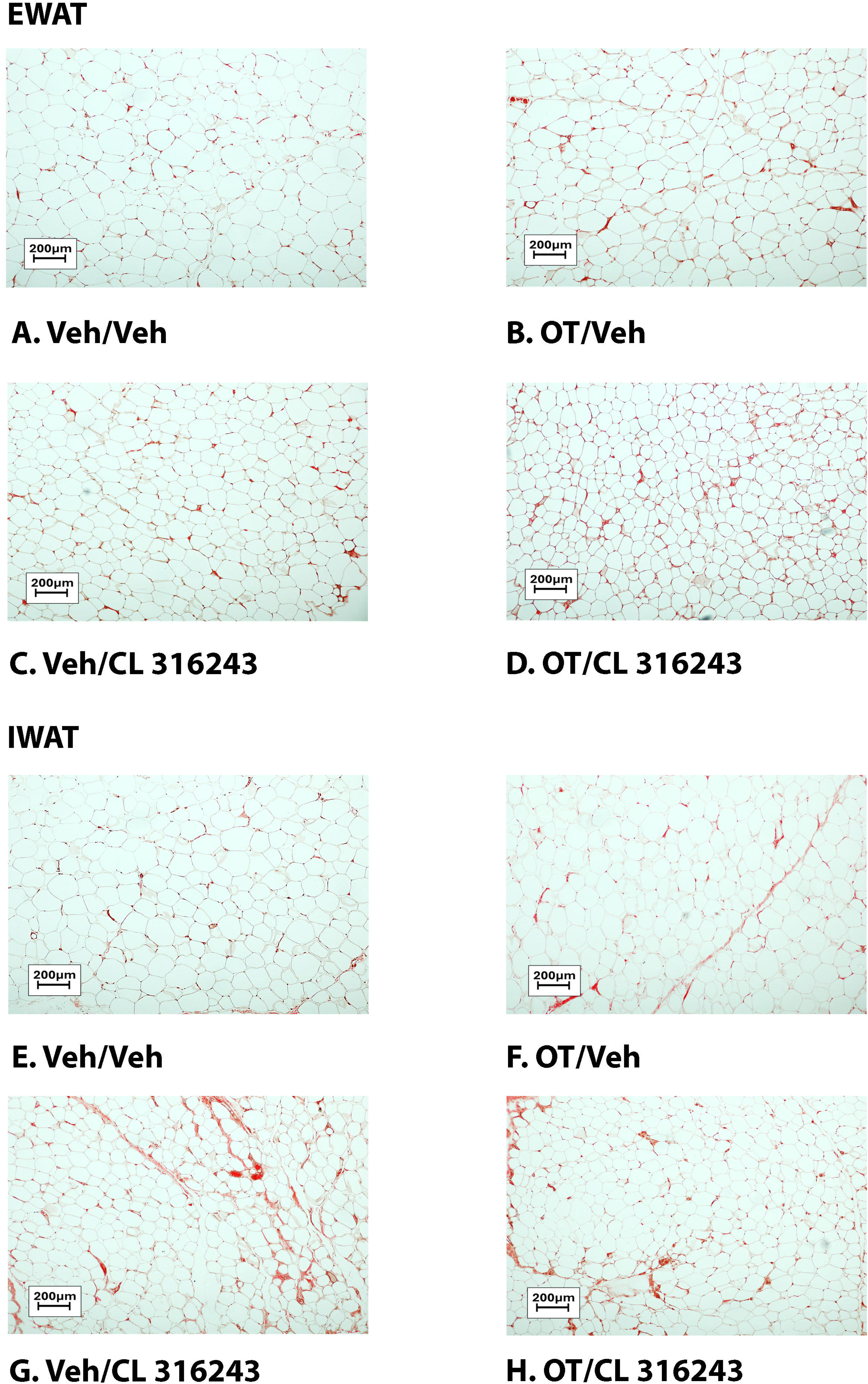
Representative image of H&E-stained section from EWAT and IWAT following chronic systemic OT infusions (50 nmol/day) and systemic beta-3 receptor agonist (CL 316243) administration (0.5 mg/kg). Images taken from fixed (4% PFA) paraffin embedded sections (5 μm) containing EWAT (A-D) or IWAT (E-H) in HFD-fed rats treated with systemic OT (50 nmol/day) or vehicle in combination with IP CL 316243 (0.5 mg/kg) or IP vehicle. A/E, Veh/Veh. B/F, OT/Veh. C/G, Veh/CL 316243. D/H, OT-CL 316243; (A–H) all visualized at 100X magnification. Images were obtained using Image Pro Plus software.

**Figure 5A-B:**
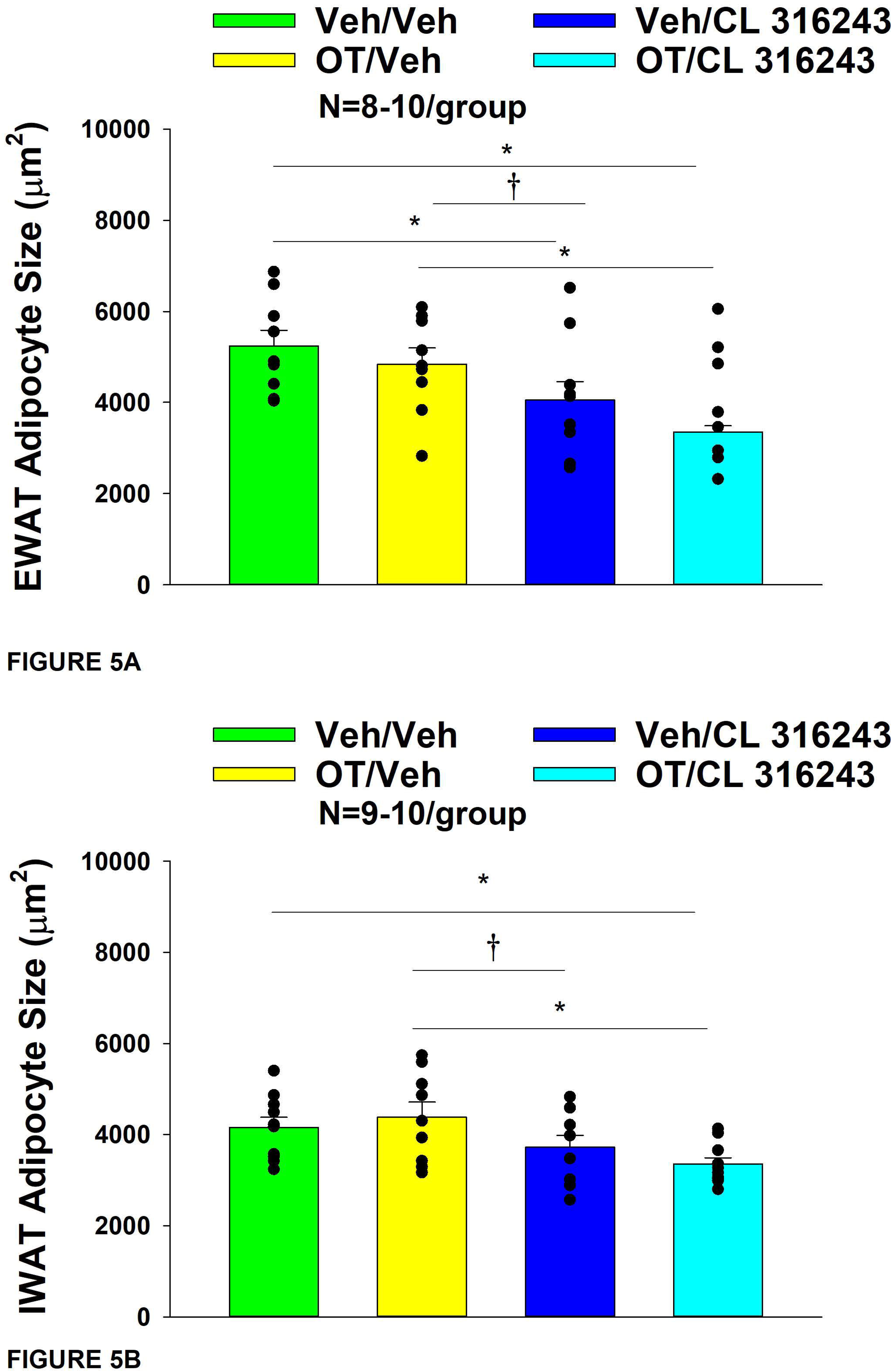
Effect of chronic systemic OT infusions (50 nmol/day) and systemic beta-3 receptor agonist (CL 316243) administration (0.5 mg/kg) on adipocyte size in EWAT and IWAT in male DIO rats. *A*, Adipocyte size (μm^2^) was measured in EWAT from rats that received chronic systemic infusion of OT (50 nmol/day) or vehicle in combination with daily CL 316243 (0.5 mg/kg) or vehicle treatment (N=9-10/group). *B*, Adipocyte size (μm^2^) was measured in IWAT from rats that received chronic systemic infusion of OT (50 nmol/day) or vehicle in combination with daily CL 316243 (0.5 mg/kg) or vehicle treatment (N=9-10/group). Data are expressed as mean ± SEM. \**P*<0.05.

### UCP-1 expression

CL 316243 alone and in combination with OT (*P*<0.05) increased UCP-1 expression in EWAT relative to VEH. In addition, CL 316243 alone also increased UCP-1 relative to OT (P<0.05). The combination of CL 316243 and OT also increased UCP-1 relative to OT treatment alone (**Figure 6A-D; Figure 7A)**, but was not different from CL 316243 alone (*P*=0.167). Similarly, CL 316243 in combination with OT (*P*<0.05) increased UCP- 1 expression in IWAT relative to VEH treatment and OT treatment (**Figure 6A-D; Figure 7A**). A subset of EWAT (N=10) and IWAT (N=3) samples were excluded in the UCP-1 expression analysis due to poor tissue quality or an insufficient amount of tissue.

**Figure 6A-D:**
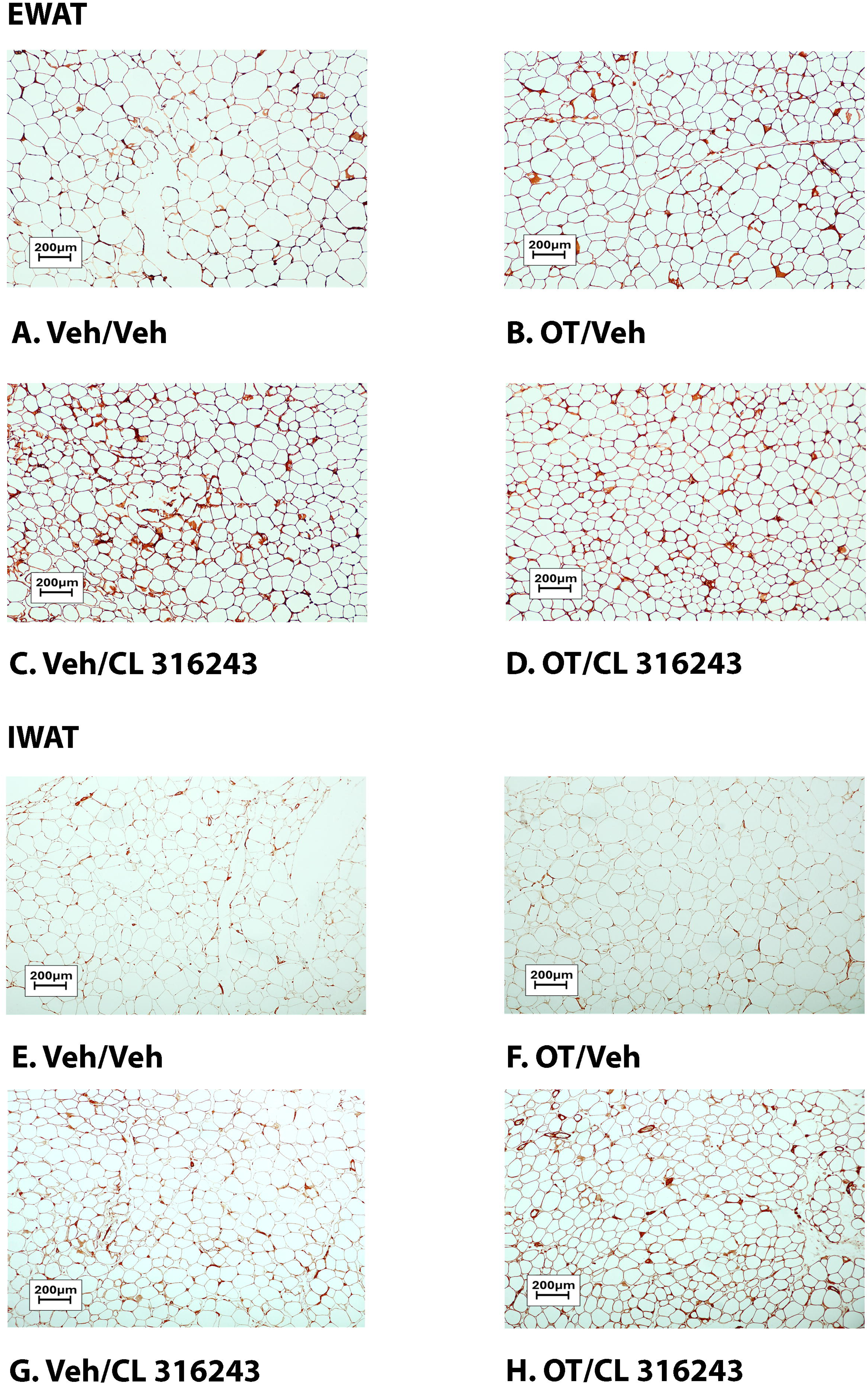
Representative image to illustrate the effect of chronic systemic OT infusions (50 nmol/day) and systemic beta-3 receptor agonist (CL 316243) administration (0.5 mg/kg) on UCP-1 content in EWAT and IWAT in male DIO rats. UCP-1 was analyzed using Image Pro Plus software. Images were taken from fixed (4% PFA) paraffin embedded sections (5 μm) containing EWAT (A-D) in HFD-fed rats treated with SC OT (50 nmol/day) or SC vehicle in combination with IP CL 316243 (0.5 mg/kg) or IP vehicle. A, Veh/Veh B, OT/Veh. C, Veh/CL 316243. D, OT/CL 316243; (A– H) all visualized at 100X magnification.

**Figure 7A-D:**
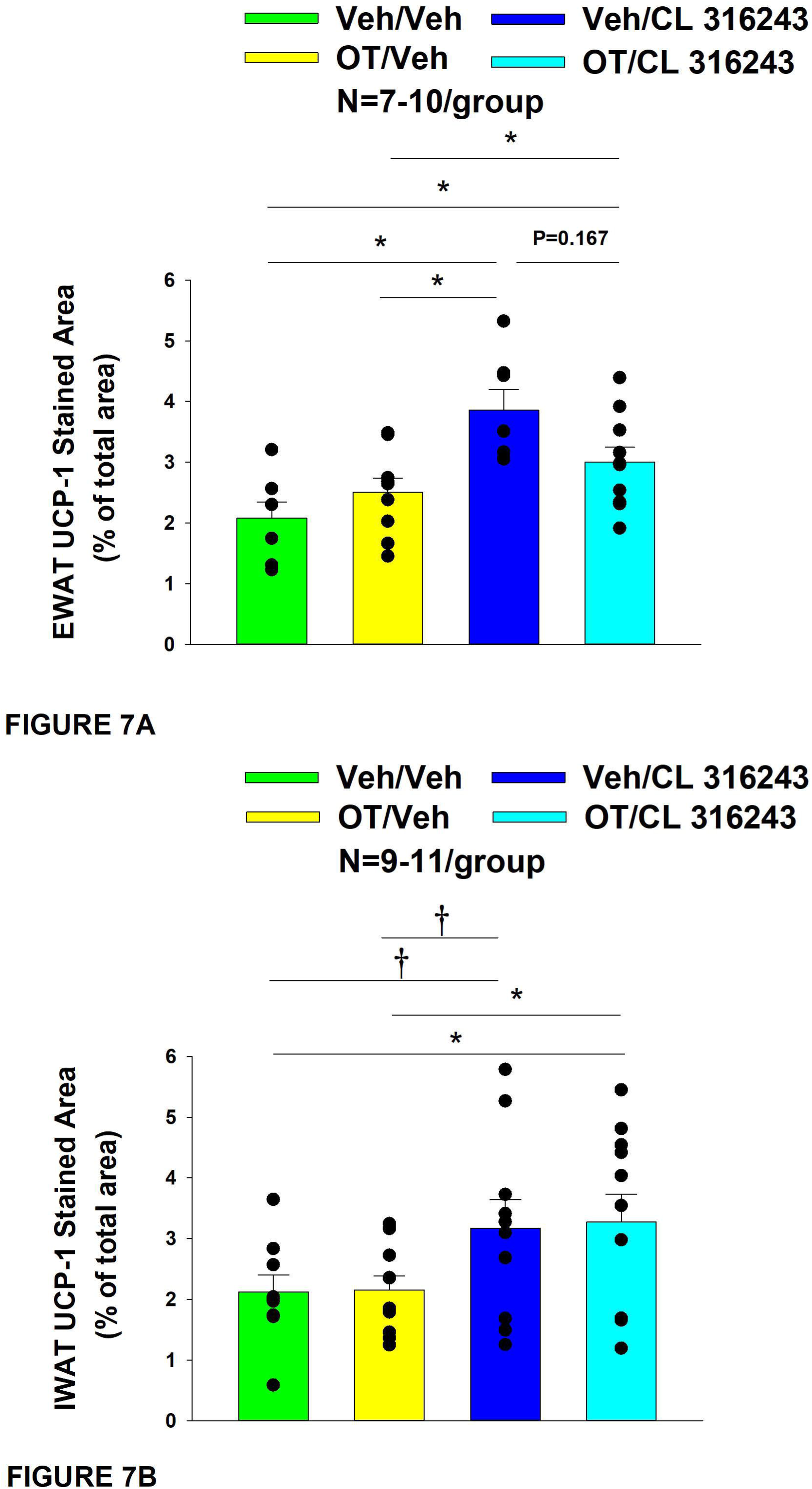
Effect of chronic systemic OT infusions (50 nmol/day) and systemic beta-3 receptor agonist (CL 316243) administration (0.5 mg/kg) on UCP-1 content in EWAT and IWAT in male DIO rats. *A*, UCP-1 expression was measured in IWAT from rats that received chronic systemic infusion of OT (50 nmol/day) or vehicle in combination with daily CL 316243 (0.5 mg/kg) or vehicle treatment (N=9-11/group). *B*, UCP-1 expression was measured in EWAT from rats that received chronic systemic infusion of OT (50 nmol/day) or vehicle in combination with daily CL 316243 (0.5 mg/kg) or vehicle treatment (N=7-10/group). Data are expressed as mean ± SEM. \**P*<0.05.

### Plasma hormone concentrations

To characterize the endocrine and metabolic effects of systemic OT (50 nmol/day) and systemic beta-3 receptor agonist (CL 316243) in DIO rats in a chronic study using as single dose of CL 316243 (**Study 3; Table 2**), we measured blood glucose levels and plasma concentrations of leptin, insulin, FGF-21, irisin, adiponectin, TC, triglycerides, and FFAs. CL 316243 alone or in combination with OT resulted in a reduction of plasma leptin relative to OT (*P*<0.05) or vehicle alone (*P*<0.05). The combination treatment was also associated with a reduction of blood glucose and insulin relative to vehicle (*P*<0.05) and OT treatment (*P*<0.05) but it was not statistically different from CL 316243 (*P*=NS). We also found that FGF-21 was reduced in response to CL 316243 and CL 316243 in combination with OT relative to vehicle and OT treatment (*P*<0.05). In addition, the combination treatment was associated with an elevation of adiponectin relative to vehicle (*P*<0.05) and OT treatment (*P*<0.05) but it was not statistically different from CL 316243 (*P*=NS). A subset of data (N=2) was excluded from the plasma hormone analysis from **Study 3** due to gross hemolysis or samples having been misplaced.

**Table 2.** Plasma measurements following systemic infusions of OT (50 nmol/day), CL 316243 (0.5 mg/kg) or OT (50 nmol/day) + CL 316243 (0.5 mg/kg) in DIO rats. Data are expressed as mean ± SEM. \**P*<0.05 vs. vehicle (N=10-12/group).

| SC | VEH/VEH | OT/VEH | VEH/CL 316243 | OT/CL 316243 |
| --- | --- | --- | --- | --- |
| Leptin (ng/mL) | 47.2 ± 6.0 | 45.9 ± 4.9 | 25.8 ± 3.4* | 22.0 ± 2.4* |
| Insulin (ng/mL) | 10.9 ± 2.3 | 11.7 ± 1.5 | 6.6 ± 1.2 | 4.3 ± 0.8* |
| FGF-21 (pg/mL) | 187.3 ± 22.5 | 163.9 ± 9.1 | 87.7 ± 8.0* | 101.5 ± 11.6* |
| Irisin (µg/mL) | 3.9 ± 0.3 | 4.5 ± 0.6 | 4.6 ± 0.4 | 4.4 ± 0.4 |
| Adiponectin (µg/mL) | 6.4 ± 0.6 | 6.3 ± 0.6 | 7.3 ± 0.6 | 8.7 ± 0.5* |
| Blood Glucose (mg/dL) | 179.3 ± 24.5 | 174.9 ± 9.3 | 150 ± 3.7 | 138 ± 4.5* |
| Triglycerides (mg/dL) | 81.5 ± 13.7 | 97.5 ± 33.1 | 39.8 ± 3.9 | 47.8 ± 3.3 |
| FFA (mEq/L) | 0.5 ± 0.1 | 0.5 ± 0.1 | 0.2 ± 0.02* | 0.3 ± 0.03<sup>a</sup> |
| Total Cholesterol (mg/dL) | 100.0 ± 6.0 | 108.1 ± 9.7 | 94.5 ± 5.1 | 107.0 ± 10.2 |

Blood was collected in 6-h fasted rats by tail vein nick (glucose) or cardiac stick
(leptin, insulin, FGF-21, adiponectin, triglycerides, FFA and total cholesterol) at 2-h
post-injection of vehicle or CL 316243 (0.5 mg/kg)
\*P<0.05 vs vehicle
N=10-12/group

### Gene expression data

We next determined the extent to which CL 316243 (0.5 mg/kg), OT (50 nmol/day), or the combination treatment increased thermogenic gene expression in IBAT, EWAT and IWAT relative to vehicle at 2-hour post-CL 316243/vehicle injections.

#### IBAT

Consistent with published findings in mice and rats, chronic CL 316243 administration elevated relative levels of the thermogenic markers *Ucp1* [52; 53; 54], *Dio2* [35; 55], *Ppargc1a* [53] and *Gpr120* [35; 53] (**Table 3A**; *P*<0.05). CL 316243 in combination with OT also elevated *Ucp1*, *Dio2, Ppargc1a* and *Gpr120* (*P*<0.05). In addition, chronic CL 316243 alone and in combination with OT reduced *Adrb3* (β3-AR) mRNA expression in IBAT (**Table 3A**; *P*<0.05).

**Table 3A-C.**
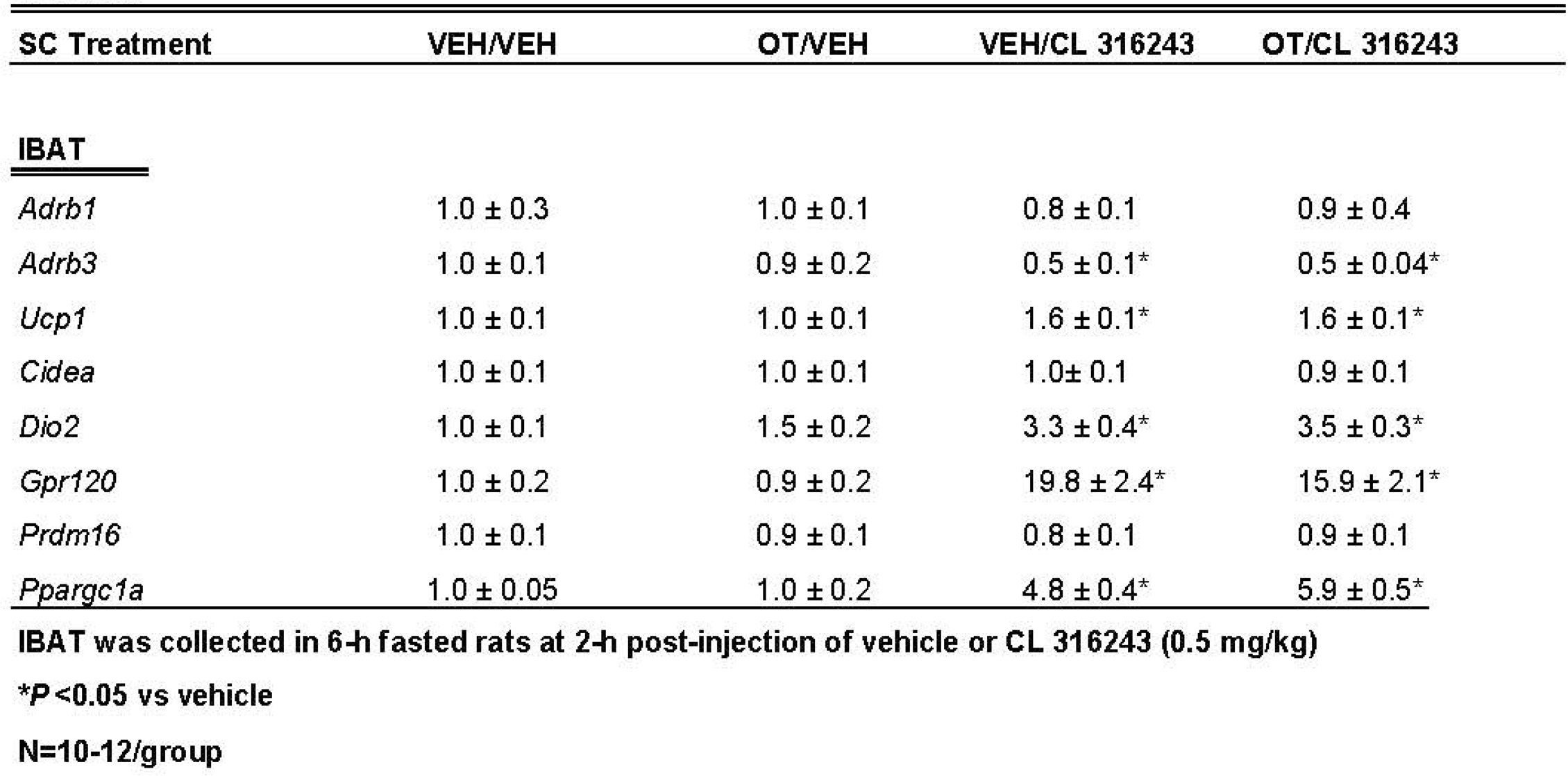
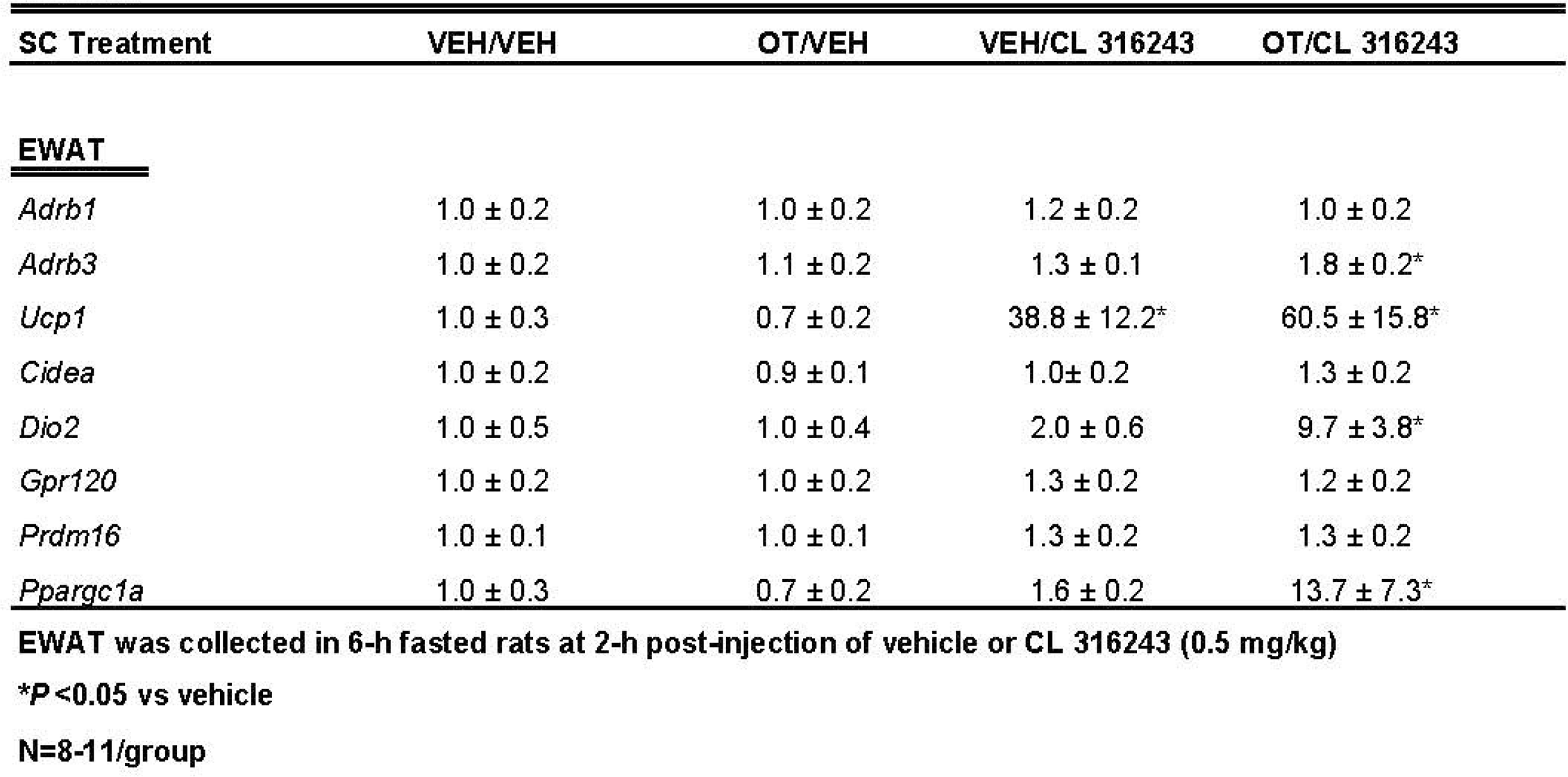
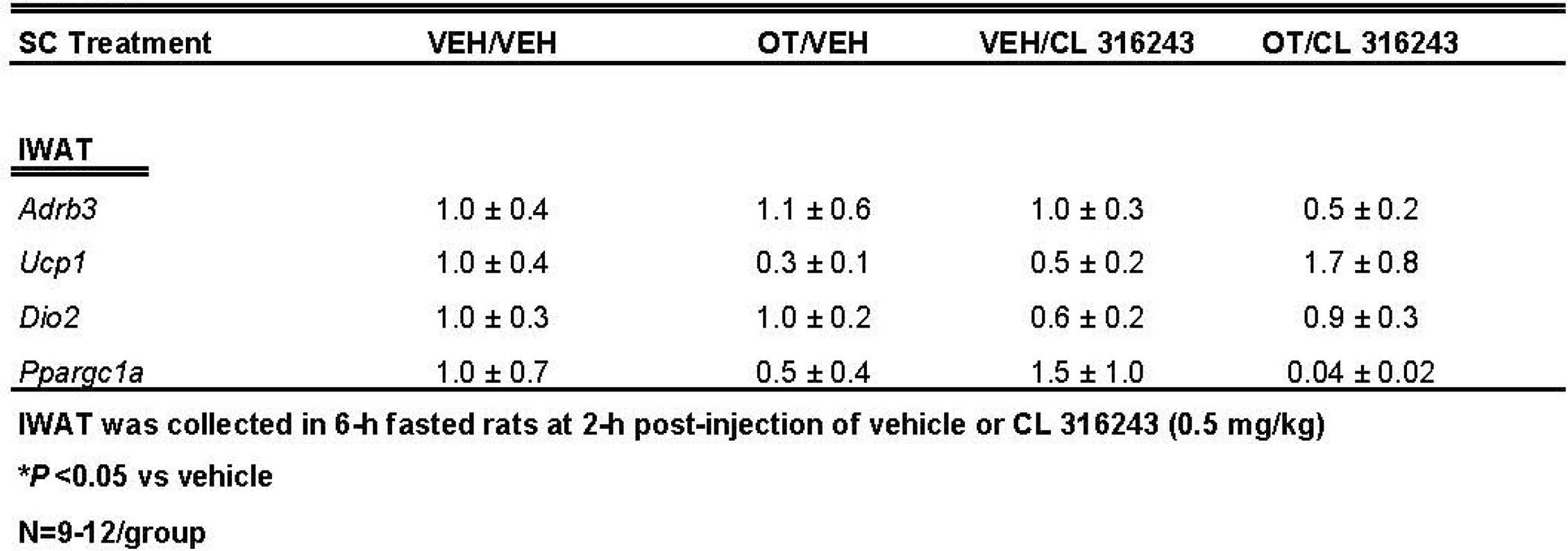
Changes in IBAT, IWAT and EWAT mRNA expression following chronic systemic OT (50 nmol/day) and systemic CL 316243 (0.5 mg/kg) treatment in male DIO rats. Data are expressed as mean ± SEM. \**P*<0.05 vs. vehicle (N=8-12/group).

#### EWAT

CL 316243 elevated relative levels of the thermogenic marker Ucp1 relative to vehicle treatment (*P*<0.05; **Table 3B**). CL 316243 in combination with OT also elevated *Ucp1*, *Dio2*, *Ppargc1a* and *Adrb3* relative to vehicle controls (*P*<0.05; **Table 3B**).

A subset of EWAT data [(N=5 (*Ucp1*); N=3 (*Dio2*); N=3 (*Ppargc1a*); N=1 (*Adrb3*), N=2 (*Cidea*)] was excluded from the EWAT gene expression analysis on account of missing samples, samples with undetectable values or statistical outliers (Grubbs’ test for outliers).

#### IWAT

There were no significant differences in the thermogenic markers *Ucp1*, *Dio2*, *Ppargc1a* or *Adrb3* in response to chronic CL 316243 alone or in combination with OT (**Table 3C**; *P*=NS).

A subset of IWAT data [(N=3 (Ucp1); N=3 (Dio2); N=1 (Ppargc1a); N=2 (Adrb3)] was excluded from the IWAT gene expression analysis on account of missing samples, samples with undetectable values or statistical outliers (Grubbs’ test for outliers).

As a functional readout of BAT thermogenesis for the gene expression analyses, we measured T_IBAT_ in response to CL 316243 alone or CL 316243 + OT during the time period that preceded tissue collection. CL 316243 alone resulted in an increase in T_IBAT_ at 0.25, 0.5, 0.75 and 1-h post-injection (**Supplemental Figure 3**; *P*<0.05). Similarly, CL 316243 + OT resulted in an increase in T_IBAT_ at 0.5-h post-injection (**Supplemental Figure 3**; *P*<0.05) and it also tended to increase T_IBAT_ at 0.75-h post-injection (**Supplemental Figure 3**; 0.05<*P*<0.1).

## Discussion

The goal of the current study was to determine the extent to which systemic OT could be used as an adjunct with the β3-AR agonist, CL 316243, to increase BAT thermogenesis and elicit weight loss in DIO rats. We hypothesized that systemic OT and beta-3 agonist (CL 316243) treatment would produce an additive effect to reduce body weight and adiposity in DIO rats by decreasing food intake and stimulating BAT thermogenesis. To test this hypothesis, we determined the effects of systemic (subcutaneous) infusions of OT (50 nmol/day) or vehicle (VEH) when combined with daily systemic (intraperitoneal) injections of CL 316243 (0.5 mg/kg) or VEH on body weight, adiposity, food intake and brown adipose tissue temperature (T_IBAT_). OT and CL 316243 monotherapy decreased body weight by 8.0±0.9% (*P*<0.05) and 8.6±0.6% (*P*<0.05), respectively, but OT in combination with CL 316243 produced more substantial weight loss (14.9±1.0%; *P*<0.05) compared to either treatment alone. These effects were associated with decreased adiposity, energy intake and elevated T_IBAT_ during the treatment period. In addition, systemic OT and CL 316243 combination therapy increased IBAT thermogenic gene expression suggesting that increased BAT thermogenesis may also contribute to these effects. The findings from the current study suggest that the effects of systemic OT and CL 316243 to reduce body weight and adiposity are additive and appear to be driven primarily by OT-elicited changes in food intake and CL 316243-elicited increases in BAT thermogenesis.

Recent findings also indicate that other agents, namely the glucagon-like peptide-1 receptor agonist, liraglutide, and the fat signal, oleoylethanolamide (OEA), act in an additive fashion with CL 316243 to reduce body weight or body weight gain. Oliveira recently reported that the combination of liraglutide and CL 316243 produce additive effects to reduce the change in body weight in a mouse model [56]. These effects appeared to be attributed to additive effects on energy intake and increased oxygen consumption in IBAT and IWAT. In addition, the combined treatment increased expression of UCP-1 in IWAT (indicative of browning). Similarly, Suarez and colleagues demonstrated that the peroxisome proliferator-activating receptor-α (PPARα) agonist and fat signal, oleoylethanolamide (OEA), act in an additive fashion with CL 316243 produced to reduce food intake and weight gain in rats [33]. These effects were associated with pronounced reductions in fat mass and increases in expression of thermogenic genes (*PPARα* and *Ucp1*) in EWAT [33]. Of interest is the finding that systemic OT and central OT can increase OEA expression in EWAT [20]. Furthermore, Deblon reported that the effectiveness of OT to decrease body weight was partially blocked in PPARα null mice [20], indicating that PPARα may partially mediate OT’s thermogenic effects in EWAT. OEA has also been found to stimulate 1) hypothalamic expression of OT mRNA [57], 2) PVN OT neurons [58], and 3) OT release within the PVN [58]. In addition, OEA also decreases food intake, in part, through OT receptor signaling [57]. Thus, it is possible that high fat diet-elicited stimulation of OEA [59] may reduce food intake, in part, through an OTR signaling and that the effects of OT to stimulate WAT thermogenesis might occur through PPARα. Additional studies that utilize adipose depot knockdown of PPARα will enable us to determine if PPARα in specific adipose depots may contribute to the ability of OT and CL 316243 to reduce body weight and adiposity.

Our findings and others raise the possibility that systemic OT could be reducing food intake and adiposity, in part, through a direct effect on peripheral OT receptors. Asker reported that OT-B12, a BBB-impermeable OT analogue, reduced food intake in rats, thus providing evidence that peripheral OTR signaling is important in the control of food intake [60]. Consistent with these findings, Iwasaki found that the ability of peripheral administration of OT to reduce food intake was attenuated in vagotomized mice [61; 62]. In addition, Brierley extended these findings and found that the effect of systemic administration of OT to suppress food intake required NTS preproglucagon neurons that receive direct synaptic input OTR-expressing vagal afferent neurons [63]. We also found that systemic administration of a non-BBB penetrant OTR antagonist, L-371257, stimulated food intake and body weight gain in rats [64]. Several studies have also found that subcutaneous infusion of OT reduced adipocyte size in 1) visceral fat in female Wistar rats that were peri-and postmenopausal [65] and 2) visceral fat in a dihydrotestosterone-elicited model of polycystic ovary syndrome in female Wistar rats [66], 3) subcutaneous fat in female ovariectomized Wistar rats [67], and 4) EWAT in male Zucker fatty rats [68]. More recent studies have confirmed that peripheral administration of long-acting OT analogues (including ASK2131 and ASK1476) also reduced both food intake and body weight [69; 70]. Together, these findings suggest that OT may also act in the periphery to decrease adipocyte size by a direct effect on OTRs found on adipose tissue [20; 21; 71]. Of translational importance is the finding that subcutaneous [20; 25; 26; 27; 28; 72] or intraperitoneal [28] administration of OT or long-acting OT analogues can recapitulate the effects of chronic CNS administration of OT on reductions of food intake and body weight. The combination treatment and CL 316243 monotherapy reduced body weight and adiposity, in part, through increased BAT thermogenesis. Both CL 316243 alone and in combination with OT elevated T_IBAT_ throughout the course of the injection study and increased IBAT thermogenic genes (UCP-1, DIO2 and Ppargc1a) and UCP-1 content in IBAT. These findings coincided with CL-316243-elicited increases in T_IBAT_ from the same animals during the time that preceded tissue collection. These findings are consistent with our previously published findings in rats [35] and other studies mice and rats that found chronic CL 316243 administration to increase the thermogenic markers DIO2 [55] and Gpr120 [52; 53; 54]. Similar to our findings, others also reported that systemic CL 316243 increased UCP-1 mRNA expression in mice [73]. We also found that the combination treatment and CL 316243 alone caused a downregulation of β3-AR mRNA expression at 2-h post-injection. This finding is consistent with we have previously reported [35] and others who have reported that both cold exposure and norepinephrine reduced β3-AR mRNA expression in mouse IBAT [74; 75] and.mouse brown adipocytes [76].

In summary, our results indicate that systemic administration of OT in combination with systemic CL 316243 treatment results in more profound reductions of body weight compared to either OT or CL 316243 alone. Moreover, the combined treatment of OT and CL 316243 stimulated BAT thermogenesis as determined by increased T_IBAT_ and thermogenic gene expression in IBAT. Together, our data support the hypothesis that systemic OT and β3-AR agonist (CL 316243) treatment produce an additive effect to reduce body weight and adiposity in DIO rats. The effects of the combined treatment on body weight and adiposity appeared to be additive and driven predominantly by OT-elicited reductions of food intake and CL-316243-elicited increases in BAT thermogenesis.

Collectively, these findings suggest that systemic OT treatment could be a viable adjunct to other anti-obesity treatment strategies. While intranasal OT has been found to reduce body weight by approximately 9.3% in a small study with limited subjects (9-11/group) [77], it was not found, however, to have any effect on body weight in a larger scale well-controlled clinical study (N=30-31/group) in which subjects were matched well for body weight, adiposity and gender [78]. Importantly, Plessow, Lawson and colleagues did find a significant effect of intranasal OT to reduce energy intake at 6-weeks post-treatment, which served as an important control. It is possible that a more extended length of treatment might have been required to take advantage of the reductions of energy intake that were not observed until the 6-week post-treatment time point. In addition, changes in dose, dosing frequency, or co-administration with Mg^2+^ [79; 80] might need to be taken into consideration in order to maximize the effects of intranasal OT on body weight in humans who are overweight or obese. Given that OT can be an effective delivery approach to reduce energy intake and elicit weight loss in several rodent models (see [16; 81; 82] for review) and obese nonhuman primates [19], it will also be important to determine if chronic systemic OT treatment can elicit weight loss when given in combination with CL 316243 at doses that are sub-threshold for producing adverse effects on heart rate or blood pressure [83]. Recent findings indicate that OT can be effective at reducing food intake and/or body weight in female rats [84] and DIO male and female mice [26], respectively. Thus, it will be important to examine if this combination treatment produces an additive effect to reduce body weight and adiposity in female DIO rodents and nonhuman primates.

## ACKNOWLEDGMENTS

The authors thank the technical support of Nishi Ivanov.

---

**Supplemental Figure 1A-B:**
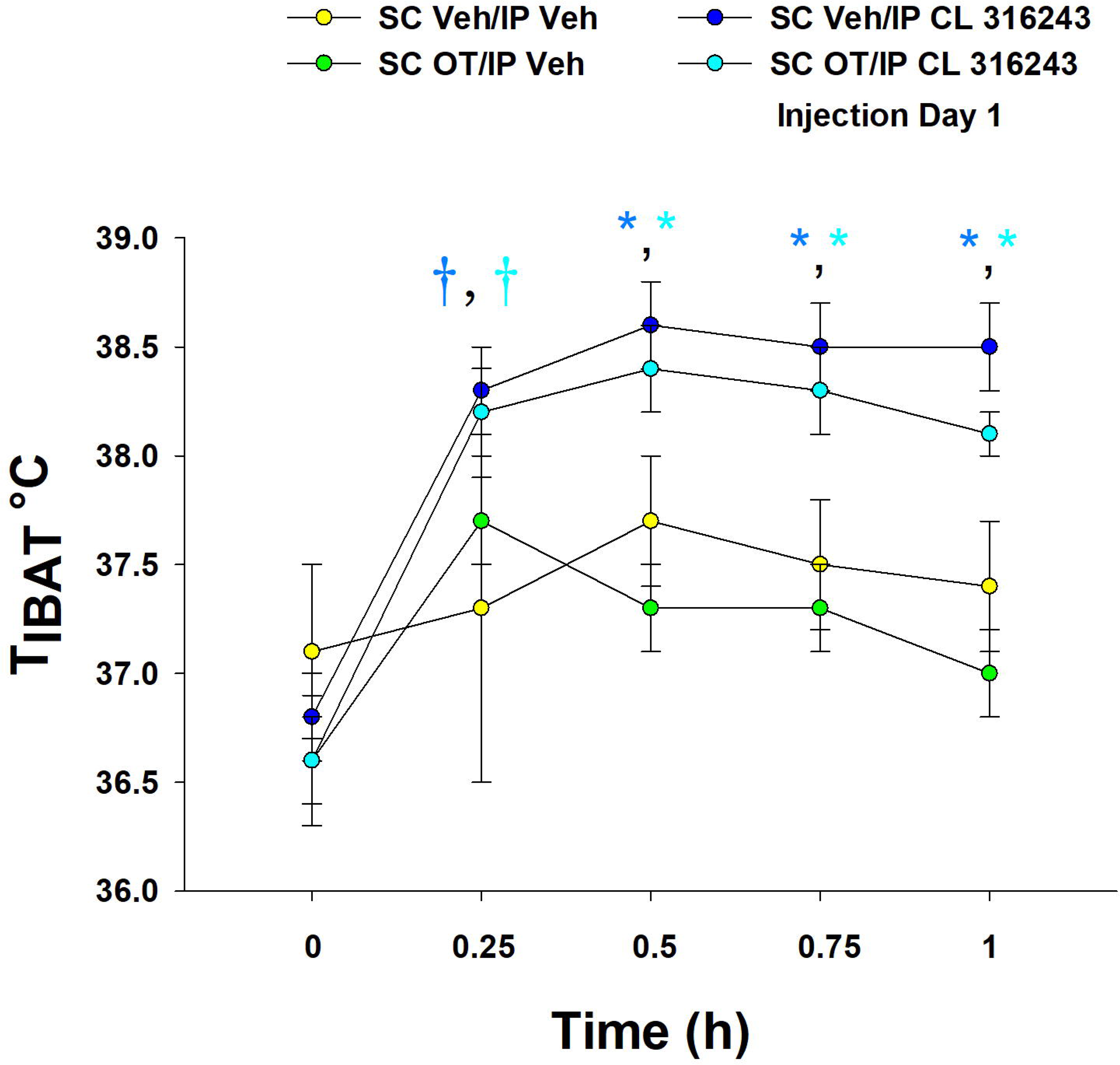
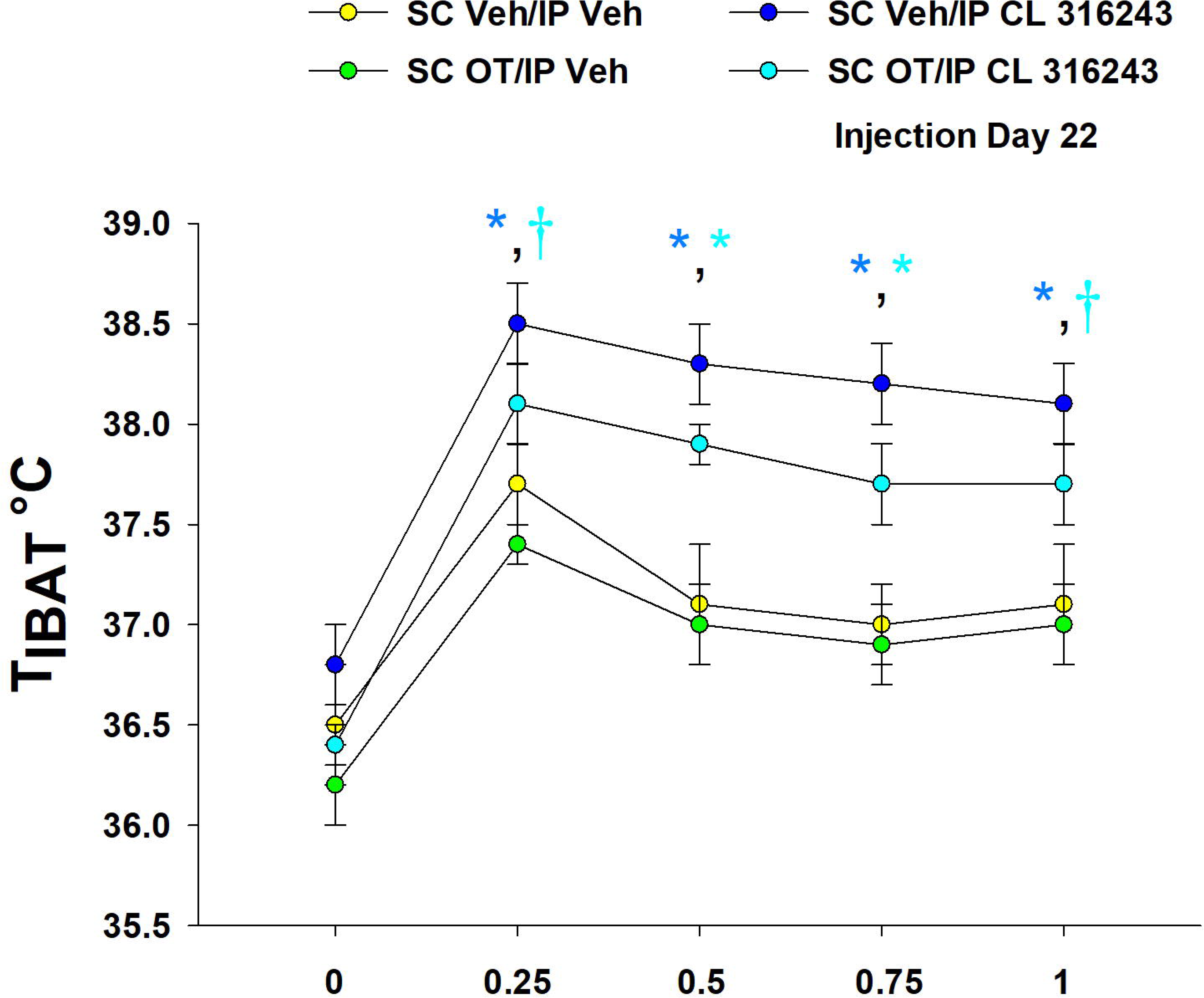
**T_IBAT_ measurements following acute systemic administration of the beta-3 receptor agonist (CL 316243) or vehicle in male DIO rats**. *A*, injection day 1 (0.5 mg/kg) and *B*, injection day 22 (0.5 mg/kg). Data are expressed as mean ± SEM. *P<0.05 vs VEH; †0.05<*P*<0.1 vs VEH.

**Supplemental Figure 2A-H:**
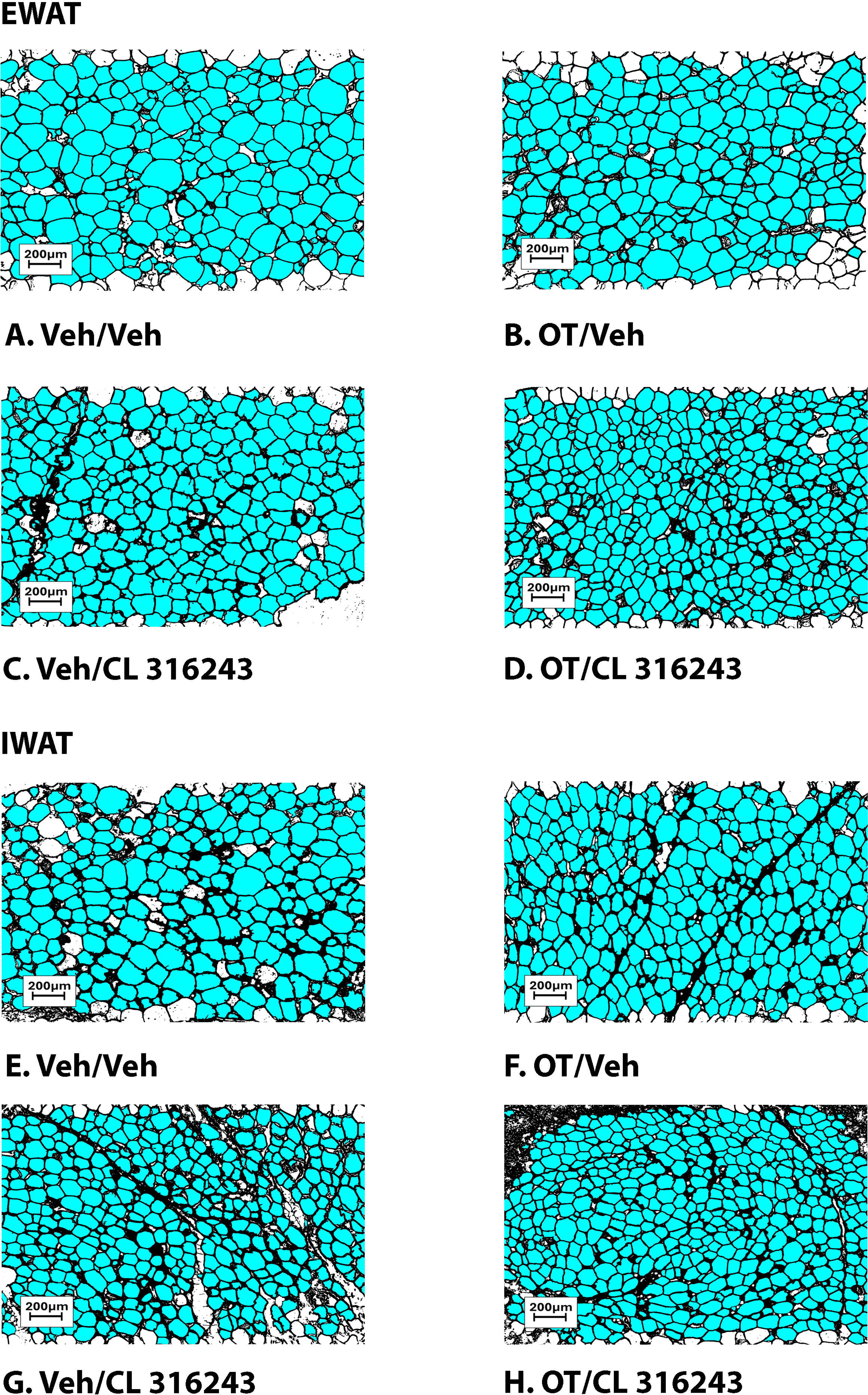
Representative image to illustrate the effect of chronic systemic OT infusions (50 nmol/day) and systemic beta-3 receptor agonist (CL 316243) administration (0.5 mg/kg) on adipocyte size in EWAT and IWAT in male DIO rats. Adipocyte size was analyzed using ImageJ. Images were taken from fixed (4% PFA) paraffin embedded sections (5 μm) containing EWAT (A-D) or IWAT (E-H) in HFD-fed rats treated with systemic OT (50 nmol/day) or vehicle in combination with IP CL 316243 (0.5 mg/kg) or IP vehicle. A/E, Veh/Veh. B/F, OT/Veh. C/G, Veh/CL 316243. D/H, OT-CL 316243; (A–H) all visualized at 100X magnification.

**Supplemental Figure 3.**
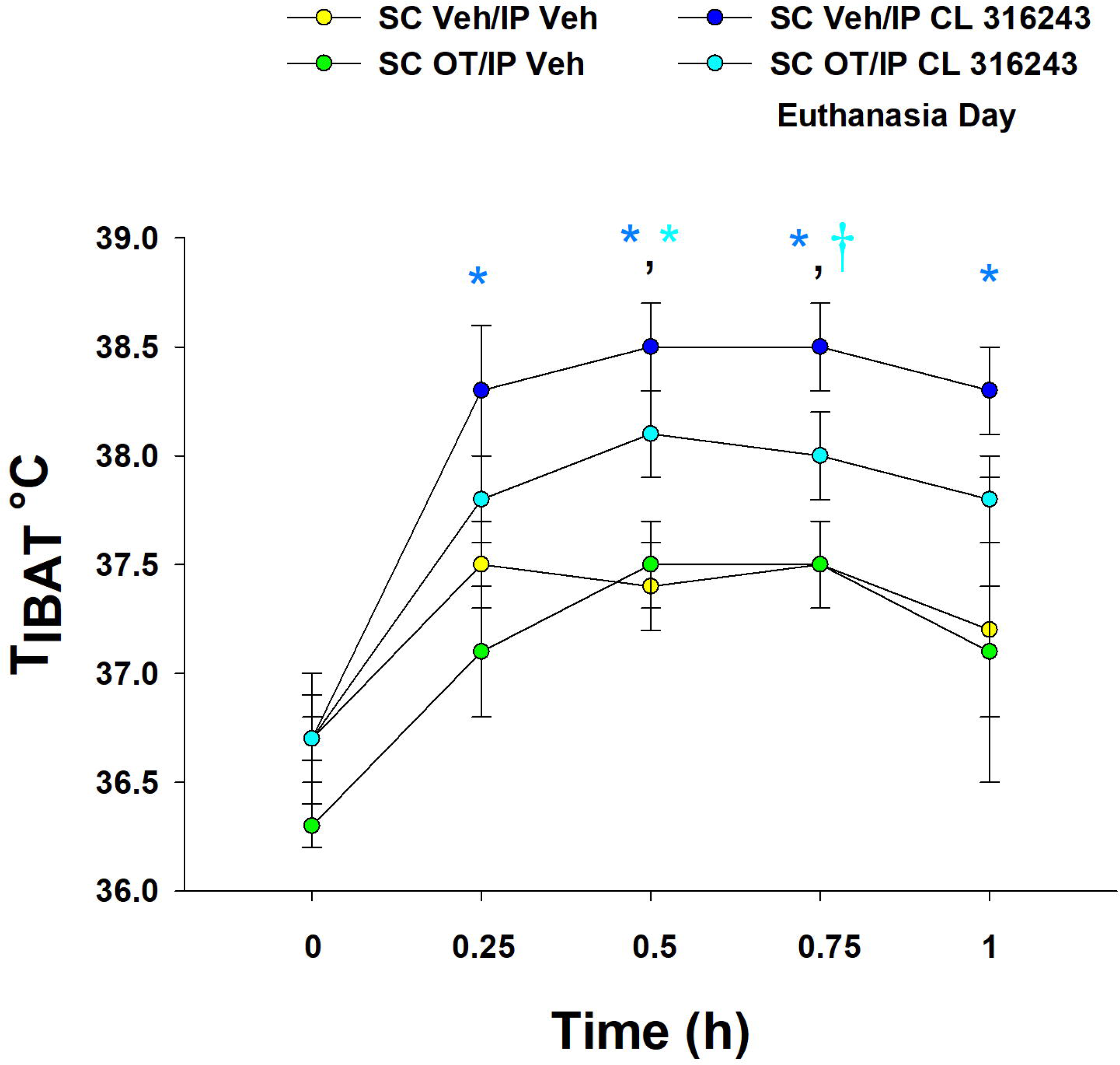
T**_I_**BAT **measurements following acute systemic administration of the beta-3 receptor agonist (CL 316243) or vehicle in male DIO rats**. *A*, injection day 23 prior to euthanasia (0.5 mg/kg), euthanasia day. Data are expressed as mean ± SEM. *P<0.05 vs VEH; †0.05<*P*<0.1 vs VEH.

